# Synergistic Combination of Cytotoxic Chemotherapy and Cyclin Dependent Kinase 4/6 Inhibitors in Biliary Tract Cancers

**DOI:** 10.1101/2020.10.26.355727

**Authors:** Mansi Arora, James M. Bogenberger, Amro M. Abdelrahman, Jennifer Yonkus, Roberto Alva-Ruiz, Jennifer L. Leiting, Xianfeng Chen, Pedro Luiz Serrano Uson Junior, Chelsae R. Dumbauld, Alexander T. Baker, Scott I. Gamb, Jan B. Egan, Yumei Zhou, Bolni Marius Nagalo, Nathalie Meurice, Eeva-Liisa Eskelinen, Marcela A. Salomao, Heidi E. Kosiorek, Esteban Braggio, Michael T. Barrett, Kenneth H. Buetow, Mohamad B. Sonbol, Aaron S. Mansfield, Lewis R. Roberts, Tanios S. Bekaii-Saab, Daniel H. Ahn, Mark J. Truty, Mitesh J. Borad

## Abstract

Biliary tract cancers (BTCs) are uncommon but highly lethal gastrointestinal malignancies. Gemcitabine/cisplatin is a standard-of-care (SOC) systemic therapy, but has a modest impact on survival and harbor toxicities including myelosuppression, nephropathy, neuropathy and ototoxicity. While BTCs are characterized by aberrations activating the cyclinD1-CDK4/6-CDKN2A-RB pathway, clinical use of CDK4/6 inhibitors as monotherapy is limited by lack of validated biomarkers, diffident pre-clinical efficacy and development of acquired drug resistance. Emerging studies have explored therapeutic strategies to enhance the anti-tumor efficacy of CDK4/6 inhibitors by combination with chemotherapy-regimens but their mechanism of action remains elusive. Here, we report *in vitro* and *in vivo* synergy in BTC models, showing enhanced efficacy, reduced toxicity and better survival with a combination comprising gemcitabine/cisplatin and CDK4/6 inhibitors. Furthermore, we demonstrated that abemaciclib monotherapy had only modest efficacy due to autophagy induced resistance. Notably, triplet-therapy was able to potentiate efficacy through elimination of the autophagic flux. Correspondingly, abemaciclib potentiated RRM1 reduction, resulting in sensitization to gemcitabine. Conclusions: As such, these data provide robust pre-clinical mechanistic evidence of synergy between gemcitabine/cisplatin and CDK4/6 inhibitors, and delineate a path forward for translation of these findings to preliminary clinical studies in advanced BTC patients.

## Introduction

Biliary tract cancers (BTCs) are uncommon but therapy-recalcitrant gastrointestinal malignancies with one of the worst prognoses among solid tumors(1). Most patients typically present with advanced, inoperable disease where standard of care (SOC) treatment is the combination of gemcitabine and cisplatin (GP) cytotoxic chemotherapy(2). Unfortunately, GP has only modest efficacy (overall response rate [ORR] ~25%, median survival ~11.7 months) and has toxicities including myelosuppression, nephrotoxicity, neurotoxicity, ototoxicity, rash and fatigue that impact quality of life, resulting in dose interruption or discontinuation (3). Almost all patients eventually experience progression of disease on first-line systemic therapy and succumb to their disease(4). As such, there is a critical clinical need to develop novel therapeutic strategies. Because BTCs represent a genomically heterogeneous tumors, targeted therapies that can be either applied ubiquitously or to more specific subsets of patients are imminently needed (5,6).

Multiple genetic alterations have been found to converge on the cyclin D1/cyclin dependent kinase (*CDK*) 4/6/ CDK inhibitor 2a (*CDKN2A*) / retinoblastoma (*RB1*) pathway in BTCs(7,8). This pathway regulates the G1-to-S cell cycle transition and its dysregulation results in uncontrolled cellular proliferation and tumorigenesis(9). Therefore, pharmacological inhibition of the cyclinD1-CDK4/6-CDKN2A-RB pathway is a rational therapeutic strategy for advanced BTC. The CDK4/6 inhibitors, palbociclib, ribociclib and abemaciclib, have shown promising pre-clinical and clinical activity and are FDA-approved in numerous solid tumors(10–12). However, palbociclib monotherapy did not demonstrate clinical activity in advanced pancreatic and biliary cancer patients with *CDKN2A* deletion and/or mutation(13). Of the three inhibitors, abemaciclib is the only one that has received approval as monotherapy for pre-treated hormone receptor positive (HR+) HER-2 negative (HER2-) metastatic breast cancer (MBC) (14). Although these three agents have not been compared within a single trial, abemaciclib response rate and tolerability are suggested to be higher than palbociclib and ribociclib (15). It has been hypothesized that putative differences in clinical activity could be attributed to the enhanced affinity of abemaciclib (~14-times greater selectivity) for *CDK4* over *CDK6*, as well as its lower myelosuppressive potential compared to palbociclib and ribociclib (16,17). Despite promising clinical advances, development of therapeutic resistance is a common characteristic of all CDK4/6 inhibitors resulting in disease progression (18,19).

Recent studies have shown that resistance to CDK4/6 inhibitors results from induction of autophagy in several solid tumors (20,21). Autophagy is a physiological stress response that maintains metabolic homeostasis and cell survival. It involves recycling cellular constituents via double-membrane vesicles called autophagosomes, which eventually fuse with late endosomes or lysosomes to form autolysosomes, facilitating degradation and generating energy for survival(22). Autophagy embroils a dynamic equilibrium of opposing roles, as a pro-survival and a pro-death mechanism. Strikingly, recent studies in solid tumor models have highlighted the importance of autophagy as a mediator of drug resistance(23,24).

Because cytotoxic chemotherapy is commonly used as standard therapy for many solid tumors, a number of preclinical and clinical studies have explored the combination of chemotherapy-regimens with CDK4/6 inhibitors. Early studies on preclinical models of pancreatic ductal adenocarcinoma (PDAC)(25), non-small cell lung cancer (NSCLC)(26), ovarian cancer (27) and triple-negative breast cancer (TNBC)(26,28–30) revealed antagonistic activity with combined cytotoxic therapy and CDK4/6 inhibition. However, recent studies demonstrated strategies for enhanced efficacy with CDK4/6 inhibitors plus chemotherapy regimens in high grade serous ovarian cancer(31,32), small-cell lung cancer (SCLC) (33,34), TNBC (35–37) and PDAC (38–40). These therapeutic strategies include protection of normal tissues, host myeloprotection and maintenance therapy for progression-free survival. The efficacy and durability outcomes of these studies have largely been attributed to the binary classification of CDK4/6-dependent and independent tumors (41). However, the predictive molecular biomarkers of the spectrum of dependence on CDK4/6 pathway are neither well understood, nor validated in clinical practice. In addition, the mechanism underlying the potential cooperative effect of CDK4/6 inhibition and cytotoxic chemotherapy remains elusive.

In this study, we performed a detailed investigation of the combination of CDK4/6 inhibition with SOC cytotoxic chemotherapy in preclinical BTC models. Our results identified the strong synergism of three FDA-approved drugs: gemcitabine, cisplatin, and abemaciclib, and elucidated the mechanistic basis for the enhanced efficacy of this triplet combination *in vitro* and *in vivo* BTC models, providing a direct path for translation for clinical investigation of these findings.

## Material and Methods

### Patient-derived xenograft studies

Animal protocols were approved by the Institutional Animal Care and Use Committee (IACUC) at the Mayo Clinic (#A00003954-18). NOD/SCID mice were obtained from Charles River Laboratory (Charles River, USA). The mice diet was LabDiet PicoLab Rodent Diet 20 (Lab Supply, Fort Worth, Texas). The mice were caged in an Innovive Disposable caging system called Innocage^®^ Mouse Pre-Bedded Corn Cob which is housed in the validated Innorack^®^ IVC Mouse 3.5 (Inno Vive, San Diego, California). Mice were housed in a 12-hour light/dark cycle with access to food and water with no fasting. Cryopreserved Patient-Derived Xenograft (PDX) tumors were implanted subcutaneously into mice as described previously(42). Tumor dimensions were monitored biweekly by manual palpation and digital caliper. When tumors reached a volume of 250-500 mm^3^, the mice were randomized according to the tumor volume into 4 groups (5 mice/group): 1) control without treatment; 2) treated with abemaciclib (45 mg/kg orally by gavage, daily); 3) treated with doublet-combination of gemcitabine and cisplatin (40 mg/kg and 4mg/kg respectively, intraperitoneal injection (IP) biweekly); 4) treated with triplet-combination of gemcitabine (40mg/kg, IP biweekly), cisplatin (4mg/kg, IP biweekly) and abemaciclib (45 mg/kg orally by gavage, daily). The dosages of gemcitabine and cisplatin were reduced to 8mg/kg and 1mg/kg respectively in PAX042. Treatment study continued for four□weeks. Tumor dimension and volume (length x width^2^) and mouse weight was recorded biweekly and compared between the four groups. Animals were euthanized on the 29th treatment day (19.5-week old mice) using cervical dislocation immediately after the cardiac puncture under anesthesia. The tumor was then harvested, measured, weighed and photographed. Immunohistochemistry (IHC) staining for Ki67, hematoxylin and eosin (H&E) and terminal deoxynucleotidyl transferase dUTP nick end labeling (TUNEL) assay were performed on FFPE tumor tissues as described previously(42).

### Statistics

Data for experiments were expressed as standard error or standard deviation of mean (median) values specified for each experiment in corresponding figure legends. Statistical tests were performed using GraphPad Prism 7.03 (San Diego, CA USA) or SAS Statistical software version 9.4 (SAS Institute, Cary, NC). The relationship between IC_50_ values and recurrent mutations across BTC cell lines was assessed by linear regression (R^2^) and Pearson correlation coefficient. Differences between groups were compared using an unpaired two-tailed t-test or non-parametric Wilcoxon rank-sum test. Tumor volume and mice weights were assessed by a mixed model using fixed effects for treatment and time (days). Survival curves were estimated using the Kaplan-Meier method and compared using the log-rank test. *P* values <0.01 were considered statistically significant.

## Results

### 1. Pharmacological inhibition of CDK4/6 in genomically characterized BTC cell lines

In order to understand the response of BTC to CDK4/6 inhibition, we employed a panel of 21 BTC cell lines, characterized by RNA sequencing and custom targeted NGS mutation profiling that reflect the genetic diversity of BTC. This cell line panel clearly recapitulates the key genomic features observed in BTC patients excluding *FGFR2* fusions but including *TP53* (67%), *IDH1* (14%), *BAP1* (5%), *PBRM1* (10%), *SMAD4* (29%), *FGFR2* (19%), *KRAS* (38%) and *CDKN2A* (62%) alterations at frequencies comparable to those observed in TCGA (**Figure 1 A, B**). mRNA sequencing analysis results showed that 60% of these cell lines lack the expression of *CDKN2A*. The gene expression levels of *CDK4, CDK6* and *CCND1* were significantly higher than *CDK2, CCNE1* and *CCNA1* suggesting that BTCs may be CDK4/6 dependent tumors **(Supplementary Figure 1)**.

**Figure 1.**
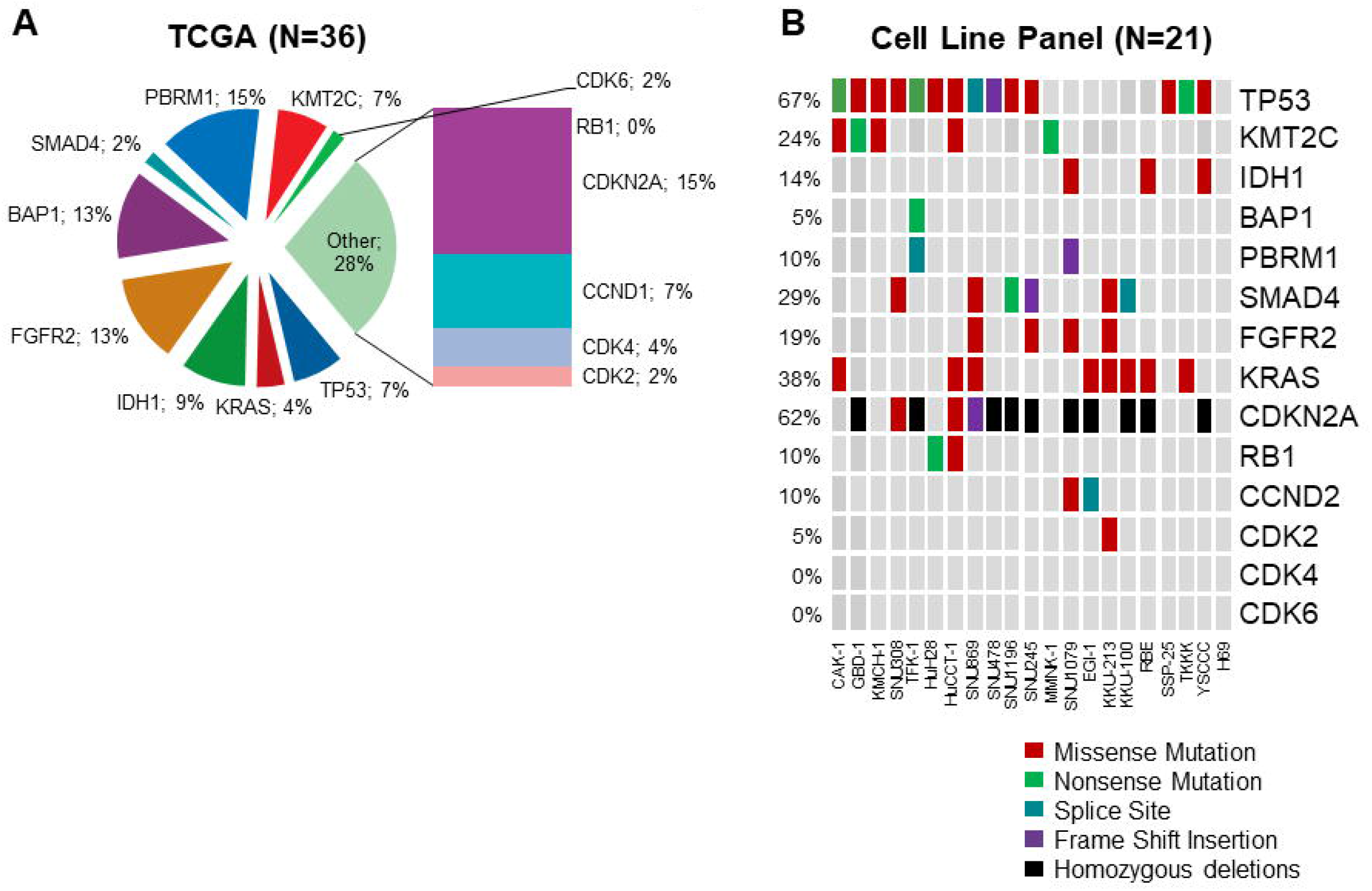
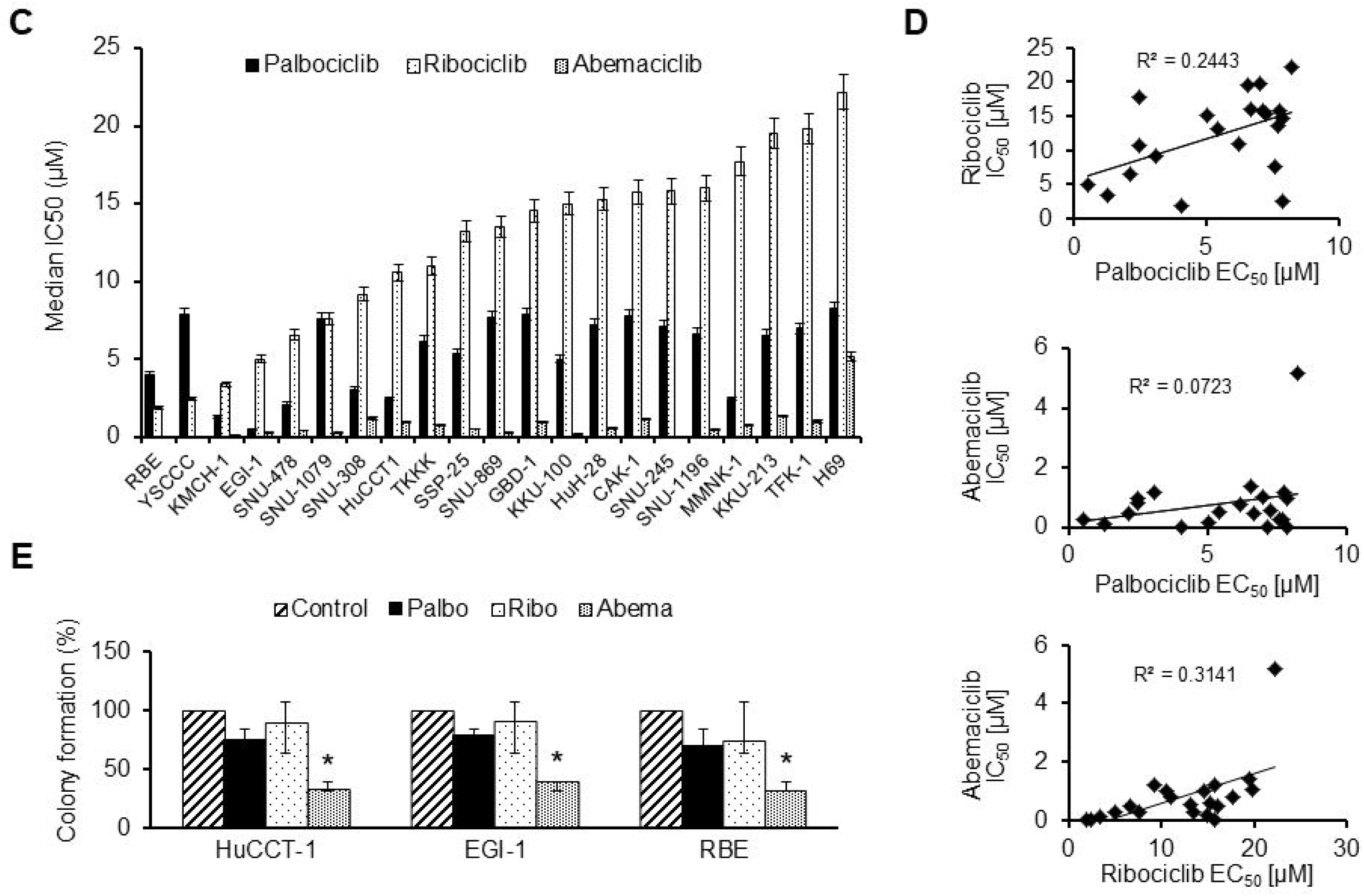
Effect of pharmacological inhibition of CDK4/6 in genomically characterized BTC cell lines. Percentage aberrations in putative driver genes shown in Cholangiocarcinoma (TCGA, The Cancer Genome Atlas; PanCancer Atlas) cohort (A) and in our panel of cell line models (B) summarized in the oncoprint (somatic missense mutations: red, nonsense mutations: green, splice site variant: blue, frame shift insertion: purple and homozygous deletion: black). (C) Mean half-maximal inhibitory concentration (IC_50_) values generated from dose response curves for palbociclib, ribociclib and abemaciclib in twenty one BTC cell lines. Palbociclib, ribociclib and abemaciclib were assessed at 10-serially-diluted doses, each dose assayed in technical quadruplicate per biological replicate. Relative cell number was determined using CellTiter-GLo signal captured on a luminescent microplate reader. Results shown are representative of three biological replicate experiments shown with median±standard deviations. (D) Linear regression analysis of palbociclib versus ribociclib; ribociclib versus abemaciclib and palbociclib versus abemaciclib IC_50_ values in BTC cell lines (N=21). (E) Three selected BTC cell lines (HuCCT-1, EGI-1 and RBE) were treated for 72 hours with 1μM of palbociclib, ribociclib and abemaciclib, prior to staining with crystal violet. Error bars represent standard deviations of mean values calculated from three independent experiments.

To evaluate sensitivity of BTC towards CDK4/6 inhibition, drug dose response (DDR) assays were performed. Treatment with palbociclib, ribociclib and abemaciclib for 144 hours significantly inhibited the viability of all BTC cell lines tested, in a dose-dependent manner, with median half-maximal inhibitory concentration (IC_50_) values of 6.58 ± 2.45 μM, 13.52 ± 5.80 μM and 0.54 ± 1.06 μM, respectively **(Figure 1C and Table 2)**. The median efficacy of abemaciclib was 12-fold or 25-fold higher than palbociclib or ribociclib, respectively. Interestingly, the IC_50_ values of palbociclib versus ribociclib (R^2^=0.2443, p=0.0227); palbociclib versus abemaciclib (R^2^=0.0723, p=0.2386); and ribociclib versus abemaciclib (R^2^=0.3141, p=0.0082), did not correlate with each other in our panel of BTC cell lines, consistent with differences in relative inhibition of CDK4 and/or CDK6, and/or variable inhibition of “off-target” kinases that may have conferred therapeutic efficacy (12,14,43) **(Figure 1D).**

To further elucidate the differences in potency of these three CDK4/6 inhibitors, crystal violet staining was used as an orthogonal assay to compare the effects of equivalent dose [1μM] of palbociclib, ribociclib and abemaciclib. We selected three cell lines (HuCCT-1, EGI-1 and RBE) that span the IC_50_ spectrum of the three CDK4/6 inhibitors. Treatment with abemaciclib for 72 hours resulted in significant reduction in colony formation (36 ± 3.3%) as compared to palbociclib (72.5 ± 4.6%) or ribociclib (89 ± 7.7%)**(Figure 1E)**. Taken together, abemaciclib was found to be most potent in BTC cell lines and was used for further evaluation in combinations.

### 2. Functional analysis of the triplet combination of abemaciclib, gemcitabine and cisplatin

Prior studies have evaluated the impact of CDK4/6 inhibitors on response to gemcitabine in different tumor types and shown evidence for either antagonistic or cooperative activity(41). To determine whether abemaciclib cooperates or antagonizes with gemcitabine/cisplatin in BTC, we performed pairwise combination DDR assays of gemcitabine and cisplatin, cisplatin and abemaciclib, and gemcitabine and abemaciclib in three selected BTC cell lines: EGI-1, RBE and HuCCT-1.

Combenefit analysis revealed the synergy levels mapped onto the combination dose-response hypersurface of viability, indicating maximum efficacy is dose dependent **(Figure 2A)**. Additivity was observed over a broad range of doses, whereas antagonism was observed at higher doses of either drug tested. For the combination of gemcitabine (0.47-1.88nM) and cisplatin (0.31-1.25μM), synergy was observed with increase in median sensitivity fold change to 2.5 ± 0.06. For maximum efficacy, a range of concentrations of abemaciclib (18.75-75.00nM) when combined with optimal doses of gemcitabine and cisplatin showed significant sensitivity gains, with median fold change sensitivity of 2.5 ± 0.09 and 2.8 ± 0.15 respectively **(Figure 2B, left panel)**. Synergism (or minimum concentration of additivity) was also confirmed using an independent scoring method by calculating the combination index (CI) values using the Chou-Talalay method **(Figure 2B, right panel)**.

**Figure 2.**
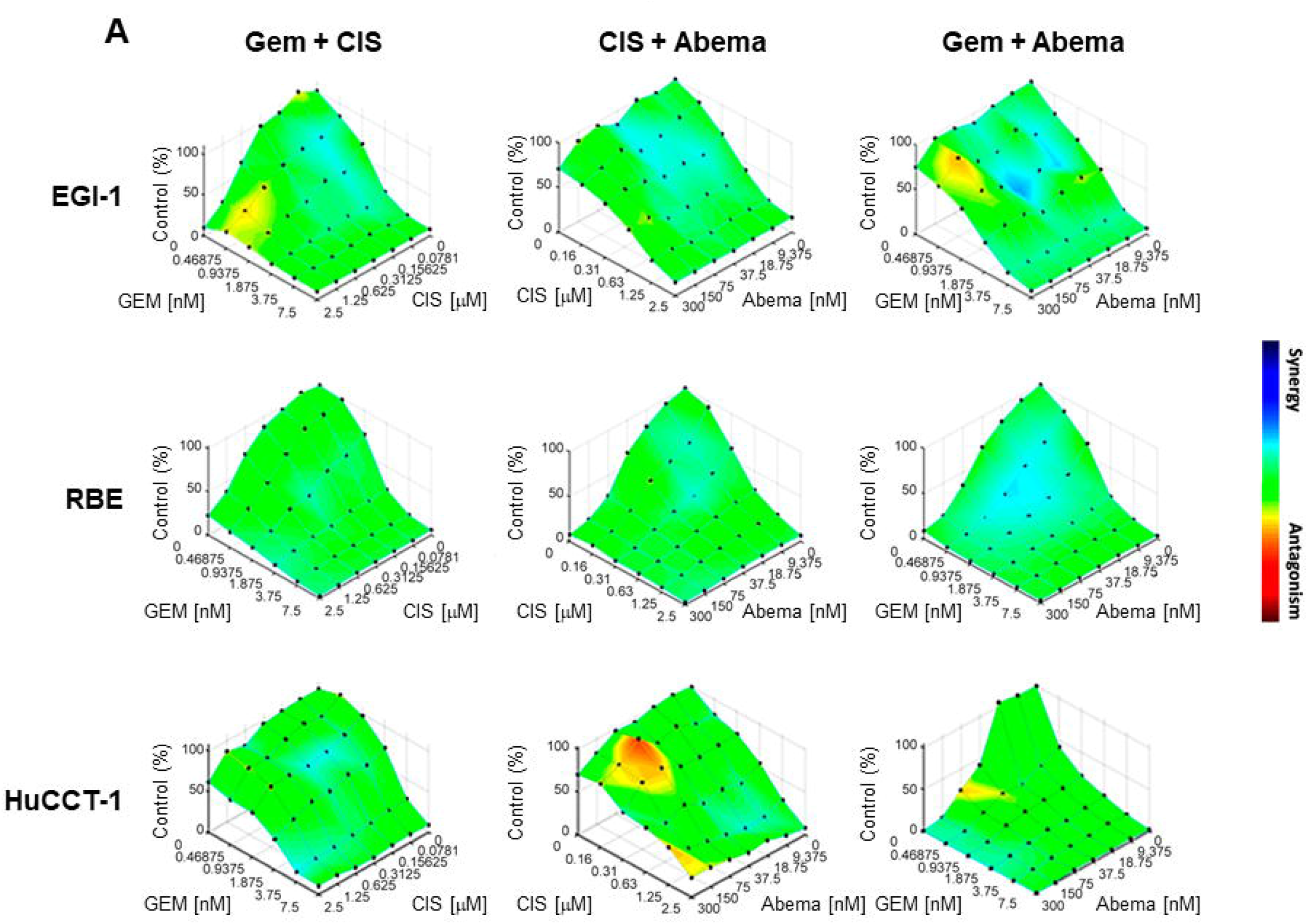
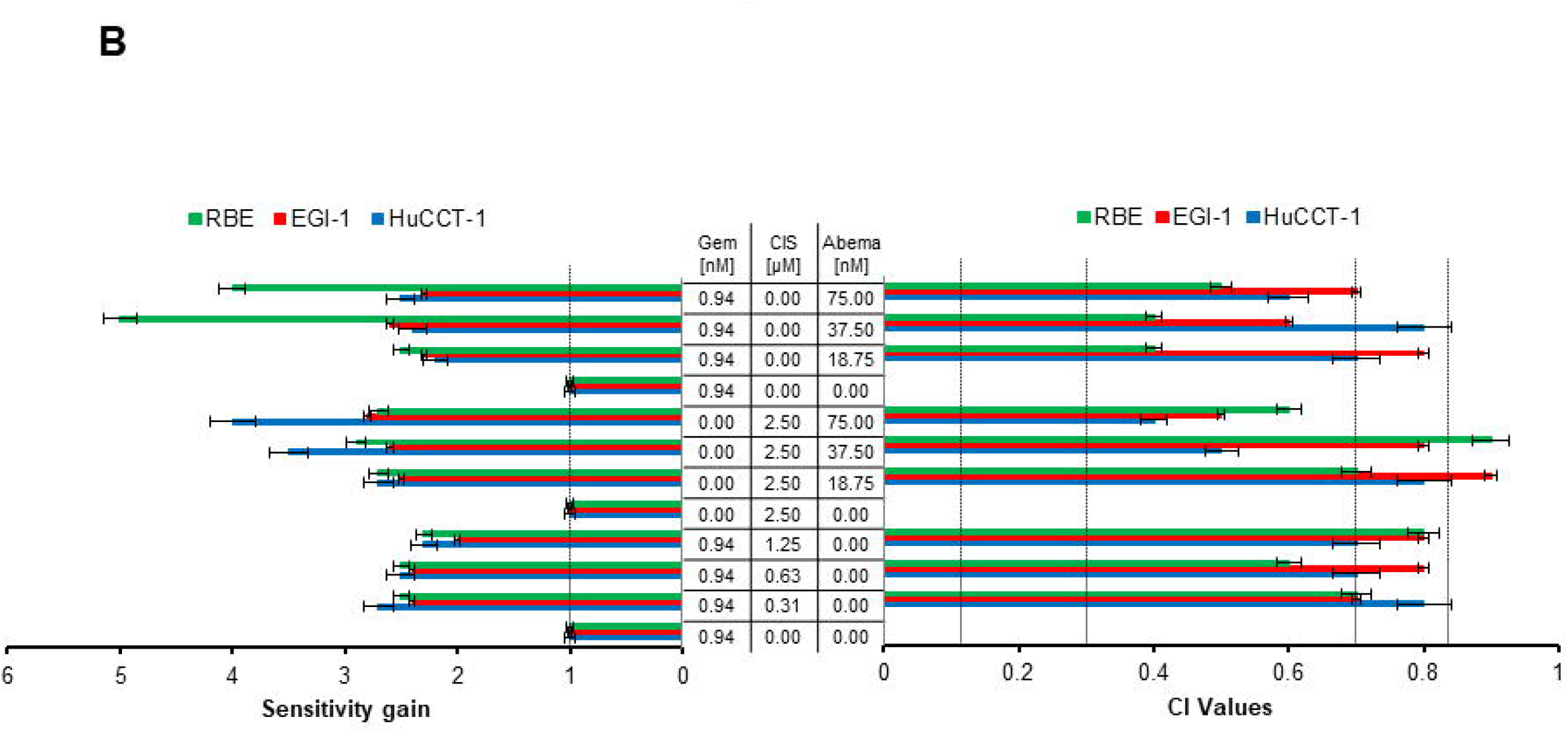
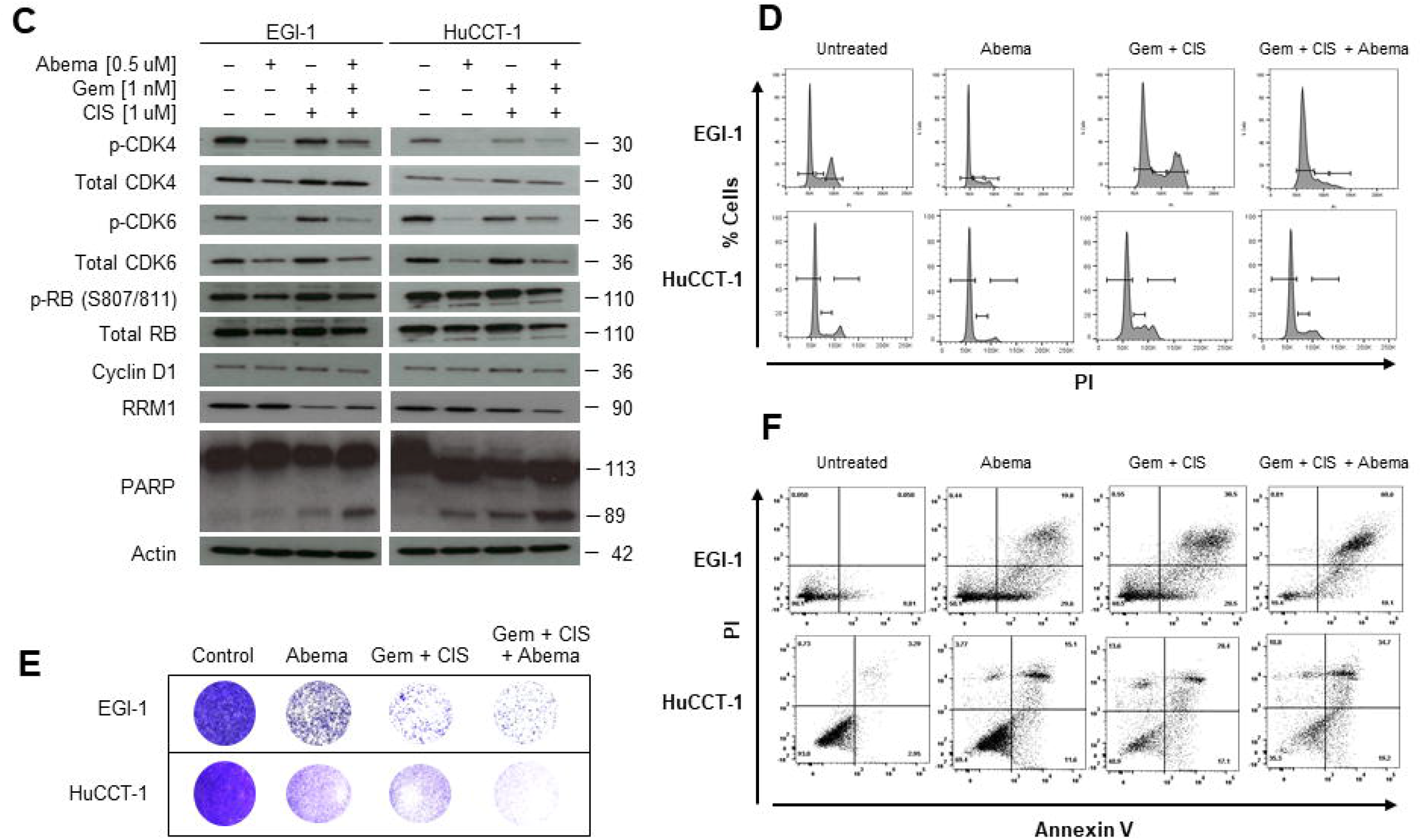
Impact of triplet combination of abemaciclib, gemcitabine and cisplatin on cellular effects of *in vitro* BTC models: (A) Combenefit plots for combination-dose response of gemcitabine (Gem) with cisplatin (CIS); CIS with Abemaciclib (Abema) and Gem and Abema in EGI-1, RBE and HuCCT-1. Synergy mapped to drug-dose response using the LOEWE model. The back wall of each plot depicts single-agent activity for each drug relative to untreated control cells (% control), while the individual data points represent the various drugdose permutations tested as indicated on the x and y axes. Drug-dose interactions are color-coded according to the key shown, with the deepest blue color indicating the strongest synergy, and deepest red color indicating the strongest antagonism. Results shown are representative of three independent experiments. (B) Fold change sensitivity gain for pair-wise combination of 3 doses of gemcitabine, cisplatin and abemaciclib (left panel) and their corresponding combination index (CI) values (right panel). Milestone CI values shown by dotted lines as >0.85 correspond to varying levels of additive to antagonism, values 0.7-0.85 correspond to moderate synergism, 0.3-0.7 represents synergism and 0.1-0.3 corresponds to strong synergism. (C) Lysates prepared from EGI-1 and HUCCT-1 cells treated for 72 hours with indicated concentrations of gemcitabine, cisplatin or abemaciclib and levels of the indicated key members of CyclinD1-CDK4/6-RB pathway proteins were measured by western blot. Images of western blots were cropped to denote the relevant band(s) for clarity; b-actin was used as a loading control. (D) Representative flow cytometric analysis plots of the indicated BTC cell lines, treated with abema [0.5μM], gemcitabine [1nM] or cisplatin [1uM] for up to 72 hours. Cellular populations at different phases of cell cycle are gated as G0-G1 (left peak), S (middle) and G2-M (right peak). Results shown are representative of three independent experiments. Cells were treated as in D prior to staining with crystal violet to assess colony formation (E) or harvesting for flow cytometry quantification of Annexin V and propidium iodide (PI) permeability as a measurement of apoptosis (F). Results shown are representative of three independent experiments.

To characterize the functional effects of the triplet combination, we investigated cyclinD1-CDK4/6-CDKN2A-Rb pathway activity by measuring phosphorylated and total protein levels of key effectors of this pathway. We observed a reduction of phospho-CDK4, phospho-CDK6, RB1 and cyclin D1 upon treatment with monotherapy abemaciclib and triplet combination as compared to the doublet combination in two selected cell lines EGI-1 and HuCCT-1 **(Figure 2C)**. Additionally, we observed a significant reduction of ribonucleotide reductase subunit M1 (RRM1) expression upon treatment with the triplet combination as compared to abemaciclib monotherapy. Next, we examined the effects of the triplet combination on the cell cycle. Cell cycle analysis indicated that abemaciclib treatment of BTC cell lines (N=3) resulted in increased percentage median population of cells at G1 phase (48.95 ± 4.4%) as compared to the untreated cells. Representative cell cycle histograms are shown for EGI-1 and HUCCT-1 **(Figure 2D)**. The triplet combination treatment resulted in a greater percentage of G1-S cells (67.4 ± 7.8%) whereas treatment with doublet combination exhibited a modest increase in percentage of G2-M cells (6.7 ± 0.3%) **(Figure 2D and Supplementary Figure 2)**.

We then assessed the inhibitory potential of the triplet combination by measuring cell counts and colony formation with clonogenic assays. Treatment with the triplet combination for 72 hours revealed percentage reduction of viable cells (75 ± 1.3%), as compared to abemaciclib monotherapy (57 ± 0.7%) or doublet-combination (70 ± 0.6%) treatment **(Supplementary figure 3A)**. Crystal violet staining of triplet-treated cells showed a significant reduction in colony formation, as compared to abemaciclib monotherapy or doublet-treated cells in all three BTC cell lines examined **(Figure 2E and Supplementary Figure 3B)**. To further characterize the cytotoxic effects of adding abemaciclib to the doublet combination, apoptotic cell death was analyzed by measuring Annexin V and PI levels by flow cytometry. We observed an increase in total apoptotic cells in response to 72-hour treatment with abemaciclib monotherapy (49%), the doublet-combination (51%), or the triplet-combination (80%) **(Figure 2F and Supplementary Figure 3C)**. Representative Annexin V versus PI plots are shown for EGI-1 and HuCCT-1 **(Figure 2F)**. We also observed a higher protein expression of cleaved PARP in triplet combination-treated cells as compared to abemaciclib monotherapy or doublet-combination treated cells **(Figure 2C)**.

### 3. Mechanistic analysis of cooperative interaction of gemcitabine-cisplatin-abemaciclib triplet combination

One of the key limitations of monotherapy CDK4/6 inhibitors is induction of therapeutic resistance by autophagy(21). We hypothesized that triplet combination might induce a therapeutic equipoise by enhancing the anti-tumor efficacy of gemcitabine/cisplatin chemotherapy and diminishing the abemaciclib-mediated autophagic evasion. Thus, we sought to investigate autophagy as the mechanism underlying the cooperative interaction of the triplet combination.

First, we assessed the presence of acidic lysosomal structures by a highly selective LysoTracker fluorescent staining. Upon corrected total cellular fluorescence (CTCF) analysis, we observed a significant decrease in LysoTracker-% fluorescence intensity in triplet-treated (19.5 ± 6.24%) cell lines as compared to the abemaciclib-monotherapy treated ones (59.5 ± 6.25%) **(Figure 3A and Supplementary figure 4A)**. Next, we evaluated the expression of LC3B-II (autophagosomal surface protein), beclin-1 (autophagy mediator) and SQSTM1 (autophagic substrate) by immunoblotting in EGI-1 and HuCCT-1. Densitometry analysis revealed increased median fold change of LC3B-II (3.1 ± 0.56) and beclin-1 (7.05 ± 0.49), concomitant with decreased SQSTM1 (0.3 ± 0.14) in abemaciclib-monotherapy treated cells **(Figure 3B)**. However, triplet combination treatment resulted in moderate decreases in median fold change of LC3B-II (1.1 ± 0.1) and beclin-1 (1.4 ± 0.7) and increased levels of SQSTM1 (1.1 ± 0.42) **(Figure 3B)**.

**Figure 3.**
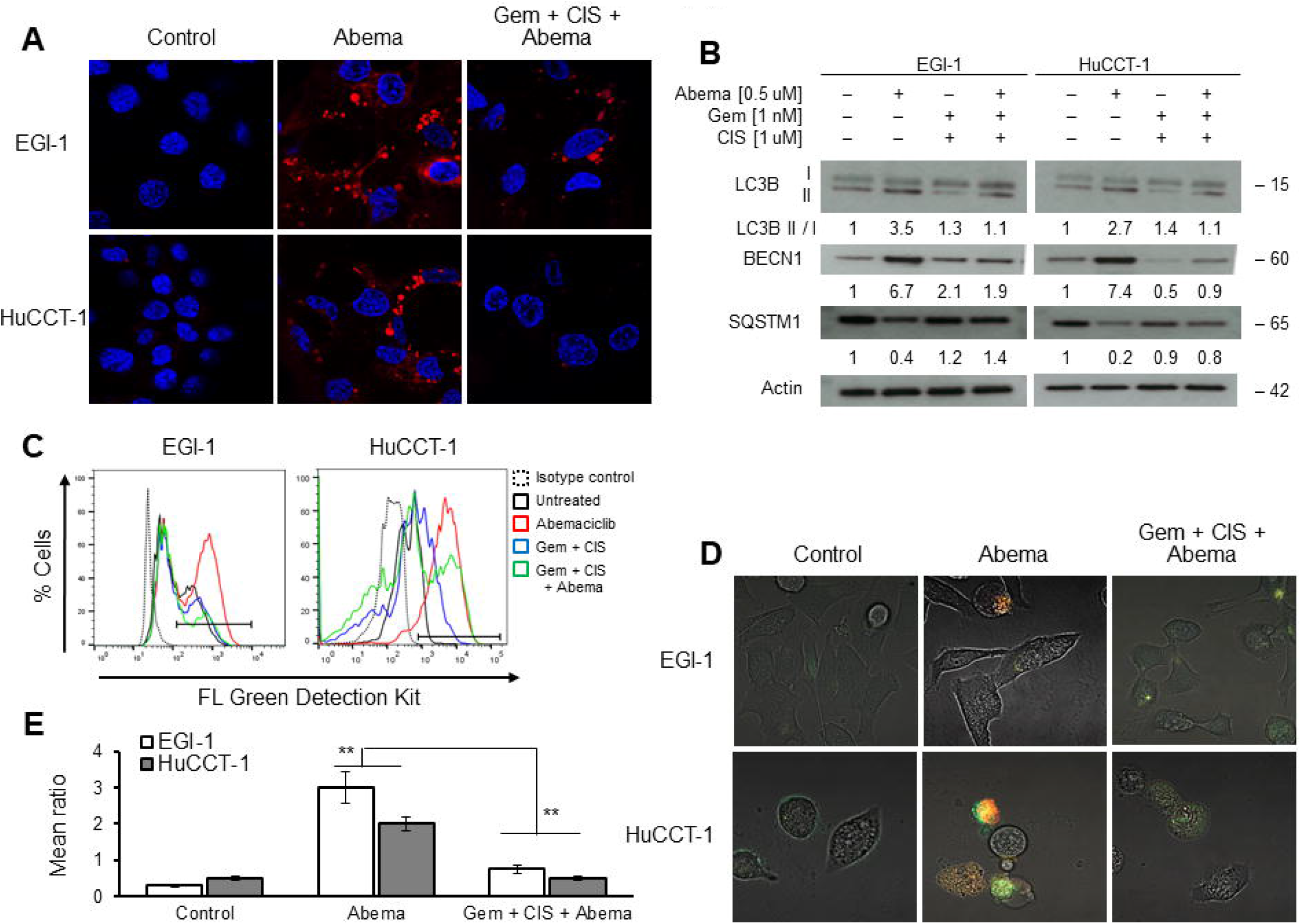
Evaluation of cellular mechanism of the cooperative effect of triplet combination. (A) Representative images of EGI-1 and HuCCT-1 cells treated with abemaciclib [0.5μM], gemcitabine [1nM] or cisplatin [1μM] for 72 hours and stained with LysoTracker deep red. Images were obtained with a confocal microscope Zeiss LSM 800 (63X) and processed with the Zen Blue software. (B) Immunoblot analysis of LC3B, BECN1 and SQSTM1 from EGI-1 and HUCCT-1 cells treated same as mentioned in A. Images of western blots were cropped to denote the relevant band(s) for clarity; b-actin was used as a loading control. (C) Representative flow cytometric analysis of EGI-1 and HUCCT-1, treated as in (A), for 72 hours and stained with green detection reagent. Untreated cells with no green detection reagent were used as isotype control to eliminate the background signal. Results shown are representative of three independent experiments. (D) Representative confocal images in EGI-1 and HUCCT-1 cells transfected with ptf-LC3 vector and treated as (A) for 48 hours. Images were obtained with a confocal microscope Zeiss LSM 800 (63X) and processed with the Zen Blue software. (E) Quantification of mean ratio of RFP-LC3 puncta and GFP-LC3 puncta. Error bars represent standard deviations of mean values calculated from three independent experiments. (**P<0.001).

Additionally, we assessed the presence of an intact autophagic flux in BTC cell lines (N=3). Firstly, we evaluated ratios of LC3B-II to LC3B-I and LC3B-II to SQSTM1 as putative markers of autophagic flux. Treatment with abemaciclib-monotherapy exhibited higher median ratios as compared to triplet combination treated cells **(Supplementary Figure 4B)**. Next, we examined the presence of an intact autophagosomal flux via a cationic-dye-based flow cytometry assay. Abemaciclib-monotherapy treated cells exhibited increased median percentage green fluorescence (87.25 ± 5.05) as compared to doublet (40.65 ± 1.16) or triplet-treated (32.65 ± 1.01) BTC cell lines **(Figure 3C and Supplementary Figure 4C)**. We then used the RFP-GFP-LC3B dual reporter assay to monitor the specific stages or flux of autophagy through LC3B protein localization. In EGI-1 and HuCCT-1, we observed a significant median increase in the RFP-puncta and GFP-RFP-puncta (expressed as mean ratio of RFP:GFP) with abemaciclib treatment (2.5 ± 0.07, p<0.001) demonstrative of an intact autophagic flux as compared to the triplet combination (0.63 ± 0.17, p<0.001) **(Figure 3D, E)**. Taken together these results demonstrate that triplet-treatment diminishes the autophagic flux associated with abemaciclib monotherapy treatment in BTC cell lines.

### 4. Evaluating anti-tumor activity of the triplet combination in cholangiocarcinoma (CCA) patient-derived xenograft models

To determine *in vivo* efficacy of the triplet combination we deployed two PDX models designated as PAX165 and PAX042, derived from patients with intrahepatic CCA. Both models were genomically characterized via aCGH and a custom targeted sequencing panel to assess copy number alterations and mutations, respectively.

PAX165 was characterized by a homozygous deletion of 9p21.3, including *CDKN2A* and *MTAP*; amplifications of 12q13.4 (*CDK2/CDK4*) and 7q21.3 (*CDK6*); a focal gain at 4q12 (*KIT* and *KDR*), and pathogenic mutations in *ATM, NCOR1* and *USP1* **(Supplementary Figure 5)**. PAX042 was also characterized by a homozygous deletion of 9p21.3 (*CDKN2A* and *MTAP*), however does not carry any other genomic alteration in the cyclinD1-CDK4/6-RB pathway.

PAX042 harbors pathogenic mutations in *ARID1A, FAT1, KRAS, PBRM1, SMAD4* and *TP53* (42). Amplification of *CDK6* has been shown to be associated with intrinsic resistance to CDK4/6 inhibitors(44). Thus, we hypothesized that PAX165 (CDK6^gain^) might be resistant to abemaciclib monotherapy as compared to PAX042 (CDK6^wt^) and that it would be interesting to see *in vivo* efficacy of the triplet combination in CDK4/6 putatively resistant and sensitive models.

In terms of tumor growth, PAX165 tumors were slower growing as compared to PAX042 **(Figure 4A, E)**. Thus, drug treatment was initiated with smaller tumors for PAX165 than for PAX042. In PAX165, abemaciclib monotherapy cohort had modest but statistically nonsignificant tumor shrinkage as compared to the control cohorts (p=0.1994). Both doublet and triplet combination treatment cohorts showed significant tumor volume differences as compared to control and abemaciclib monotherapy treatment cohorts in PAX165 (p=0.0184 for doublet, and p=0.0137 for triplet-combination). However, doublet and triplet treatment cohorts did not demonstrate any tumor volume differences when compared to each other, suggesting the need to optimize doses of gemcitabine and cisplatin **(Figure 4A)**. Similarly the body weight of mice in the abemaciclib monotherapy cohort were marginally less than the control cohort, but differences were not statistically significant (p=0.2279). The weight of mice from the doublet cohort were significantly lower than that of triplet, abemaciclib-monotherapy or control cohorts (p=0.0043 for doublet and p=0.0418 for triplet-treated) **(Figure 4B)**. Survival curves yielded superior survival in the triplet combination cohort, in contrast to the doublet or abemaciclib monotherapy cohorts **(Figure 4C, D)**.

**Figure 4.**
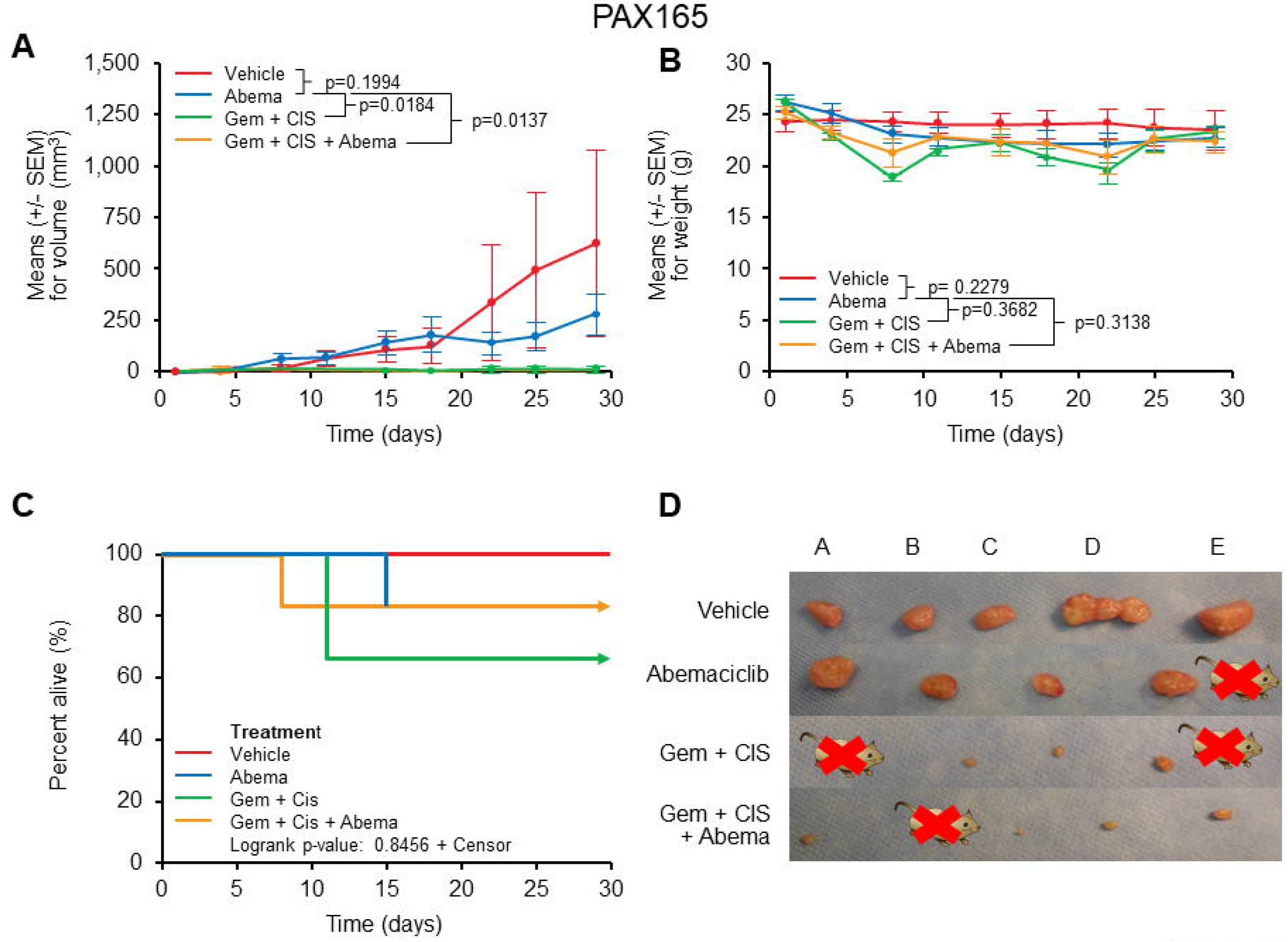
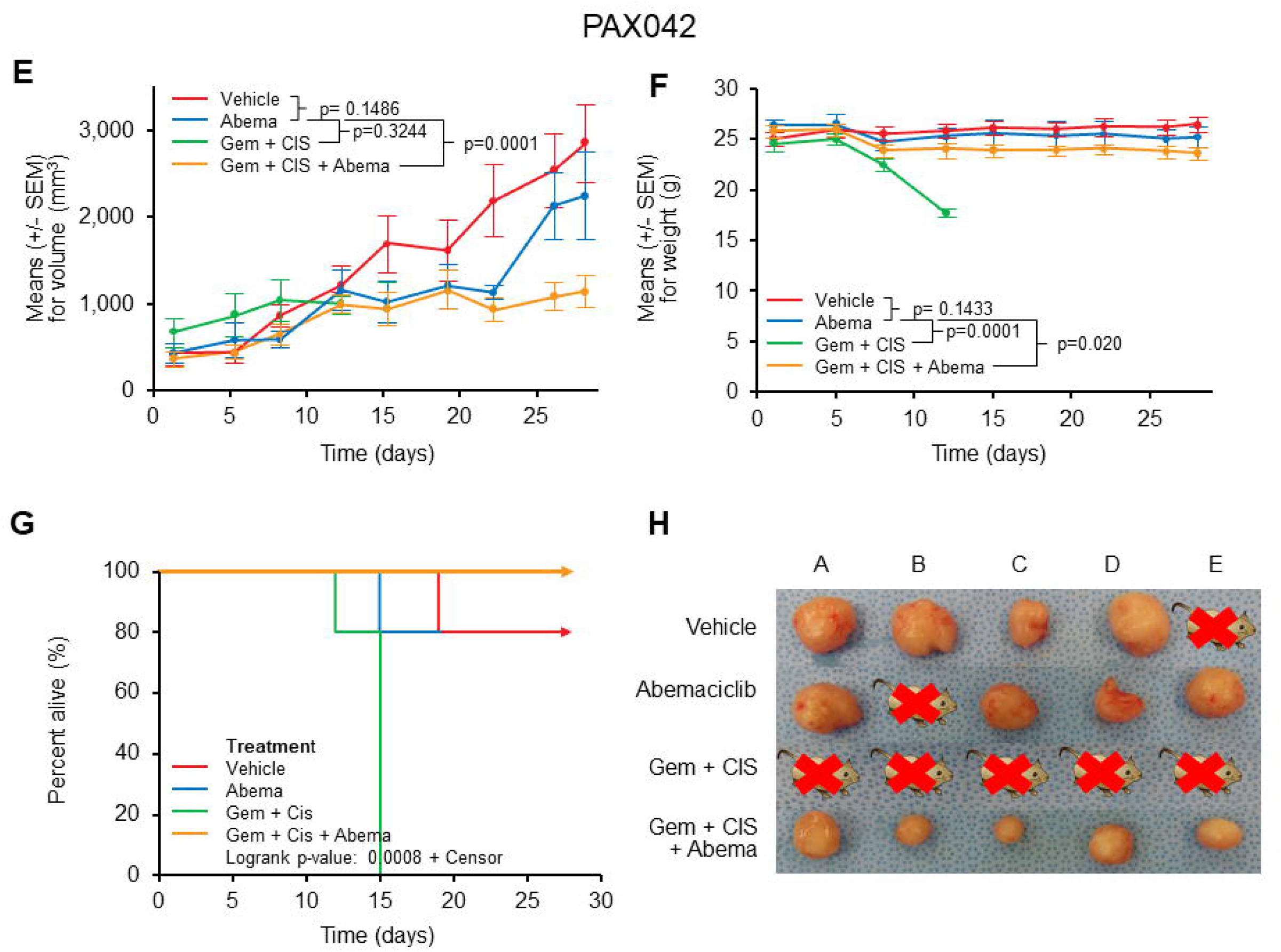
*In vivo* efficacy of triplet combination assessed in cholangiocarcinoma PDX models. The indicated PDX models were randomized for treatment with vehicle, abemaciclib (abema), doublet-combination of gemcitabine with cisplatin (gem + cis) and triplet-combination of gem, cis and abema (gem + cis + abema) when the tumor volume reached 250-500mm^3^ for four weeks. Tumor volumes and mouse weights were recorded biweekly. For PAX 165 tumor volume (A), mouse weight (B), Kaplan-Meier survival curve with death and tumors exceeding 1000mm^3^ as end point (C) representative images of the tumors that were excised from the mice at the end of the treatments (D) were compared across the four groups. For PAX042, tumor volume (E), mouse weight (F), Kaplan-Meier survival curve with death and tumors exceeding 1000mm^3^ as end point (G) representative images of the tumors that were excised from the mice at the end of the treatment (H) were compared across three groups. Animals were euthanized on the 29th treatment day. Data represented as mean ± s.e.m. n=5 for each group. (***P<0.0001, **P<0.001, *P<0.01).

Since the anti-tumor activity of doublet and triplet-combination treatment cohorts did not show any difference in PAX165, we reduced the dosages of gemcitabine to one fifth and cisplatin to one fourth for investigation of PAX042 (8mg/kg gemcitabine and 1mg/kg cisplatin), to discern the contribution of adding abemaciclib to the doublet. In PAX042, we did not observe any statistically significant tumor volume differences in abemaciclib-treated cohorts as compared to the control cohorts (P=0.1486), contradicting our original hypothesis of PAX042 being a CDK4/6 sensitive model. Surprisingly, all mice died in the doublet treated cohort at day 15. In the triplet-treated cohort, tumor volume differences were significantly greater as compared to the abemaciclib monotherapy and control cohorts (p=0.0001) **(Figure 4E)**. The bodyweight of mice in abemaciclib-treated cohort were similar to that of the control cohort (p=0.1443). In contrast, doublet-treated cohort showed a gradual drop of bodyweight by day 15 when all the mice died (p=0.0001). The triplet-treated cohort mice showed slight drop in body weights but it was not statistically significant (p=0.0200) indicating that abemaciclib mitigates the toxicity of doublet-chemotherapy treated mice **(Figure 4F)**. Survival curves suggested a better survival for the triplet treated cohort (with zero mice deaths) as compared to the doublet-treated (all mice died) or abemaciclib monotherapy (one mice died) treated cohorts **(Figure 4G, H)**. Taken together, these results clearly demonstrate that addition of abemaciclib renders a survival benefit to the doublet-chemotherapy treated mice. However, additional studies with higher statistical power are required to make definitive conclusions.

### 5. Evaluation of pharmacodynamic and functional markers of triplet-combination in *in vivo* BTC models

Tumor staining of proliferative marker Ki-67 has been correlated with CDK4/6 inhibitor response in multiple preclinical studies(45). We observed a significantly reduced Ki67^+^ staining in tumor xenograft tissues from triplet-treated cohorts. In the PAX165 model, a 44% decrease in Ki67^+^ cellular proliferation in the abemaciclib-treated cohort (p=0.0001), 82% in doublet-treated (p=0.0003), whereas 80% decrease was observed in triplet-treated (p=0.0001) as compared to the control cohort tumor tissues **(Figure 5A, top panel)**. Similarly, in PAX042, we observed a 42% decrease in Ki67+ cellular proliferation in abemaciclib treated cohort (p=0.0009) and a 67% decrease in triplet-treated (p=0.0002) as compared to the control cohort **(Figure 5B, top panel)**. This suggests that the triplet combination treatment reduces the proliferative capacity of CCA PDX models.

**Figure 5.**
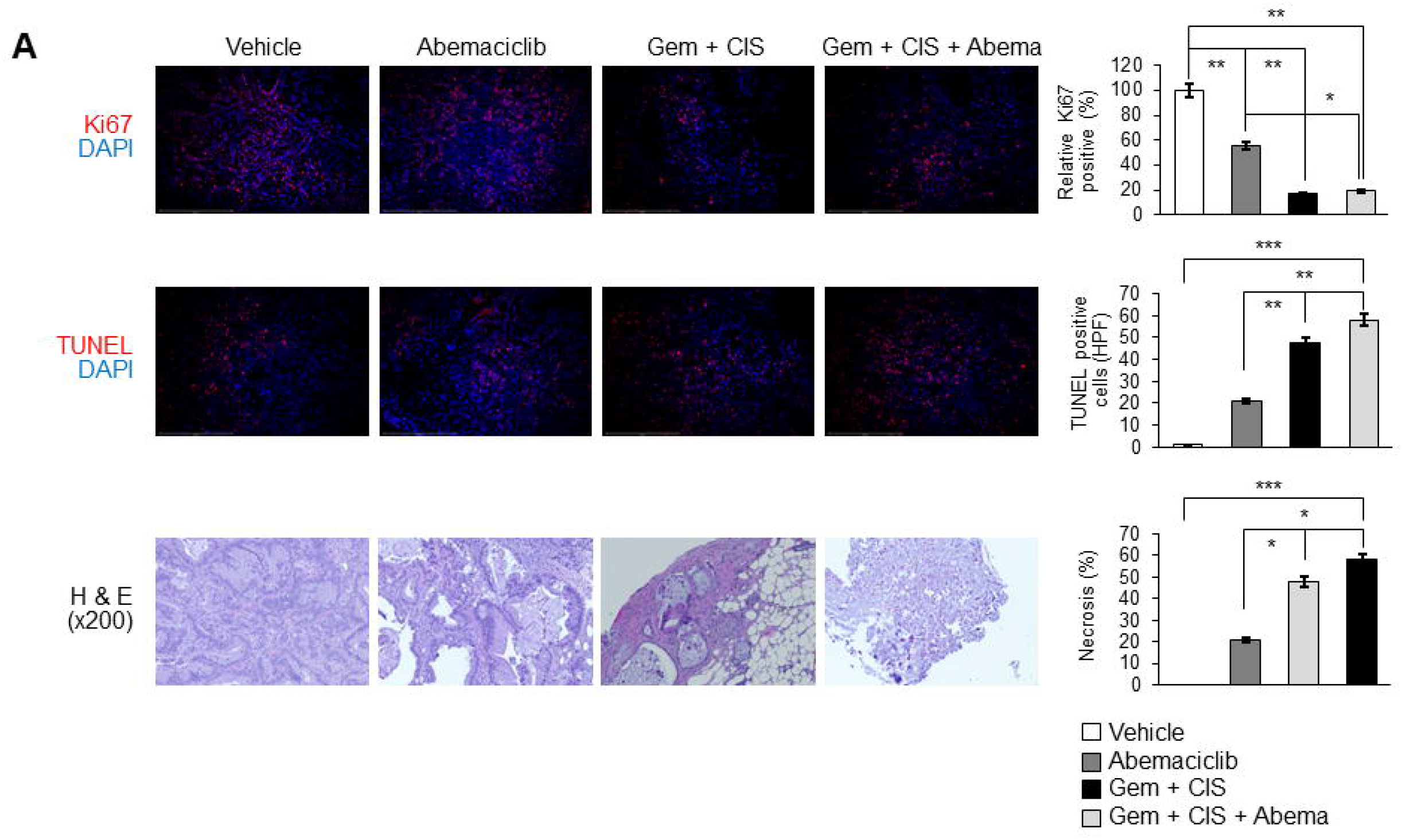
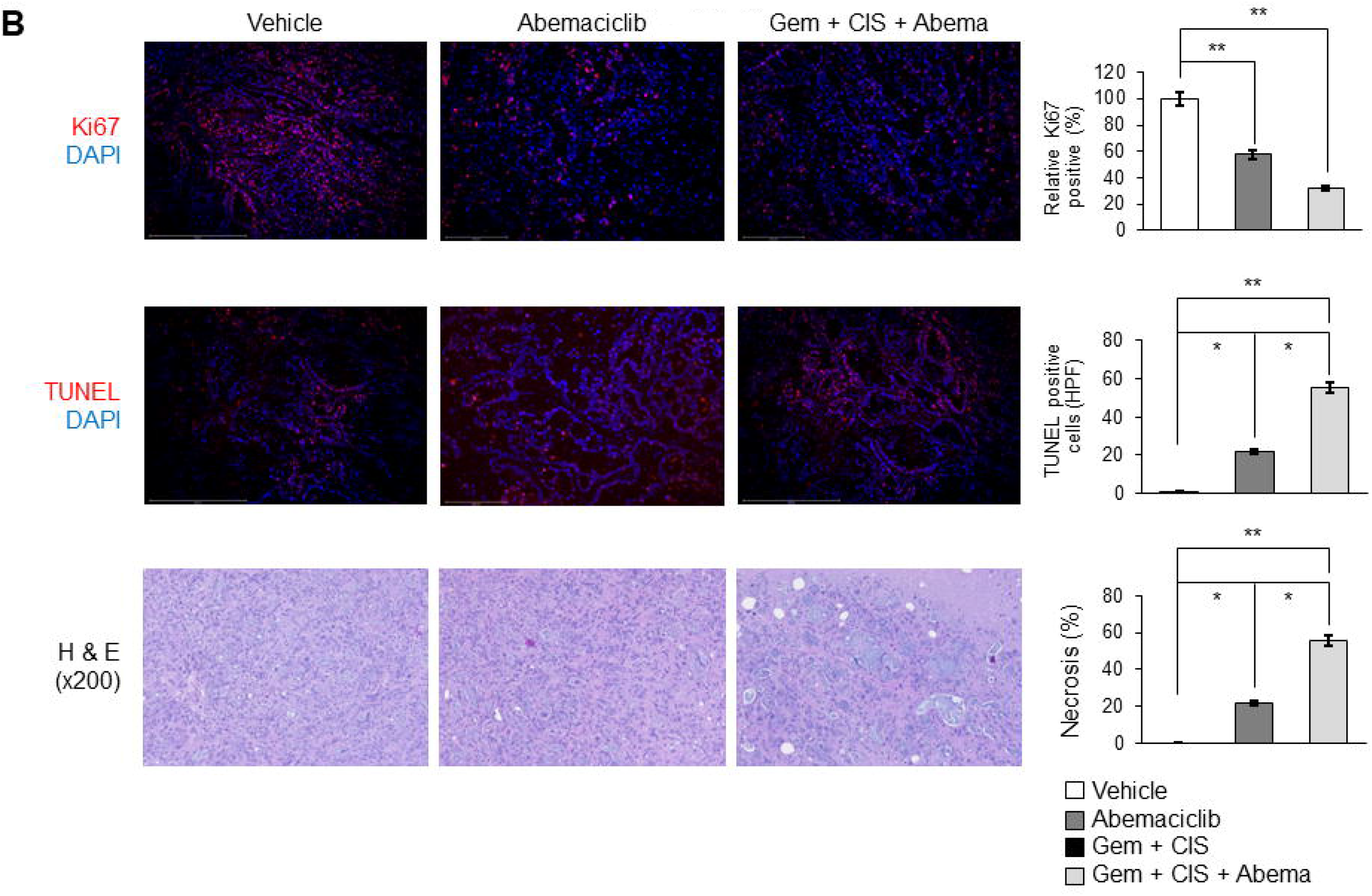
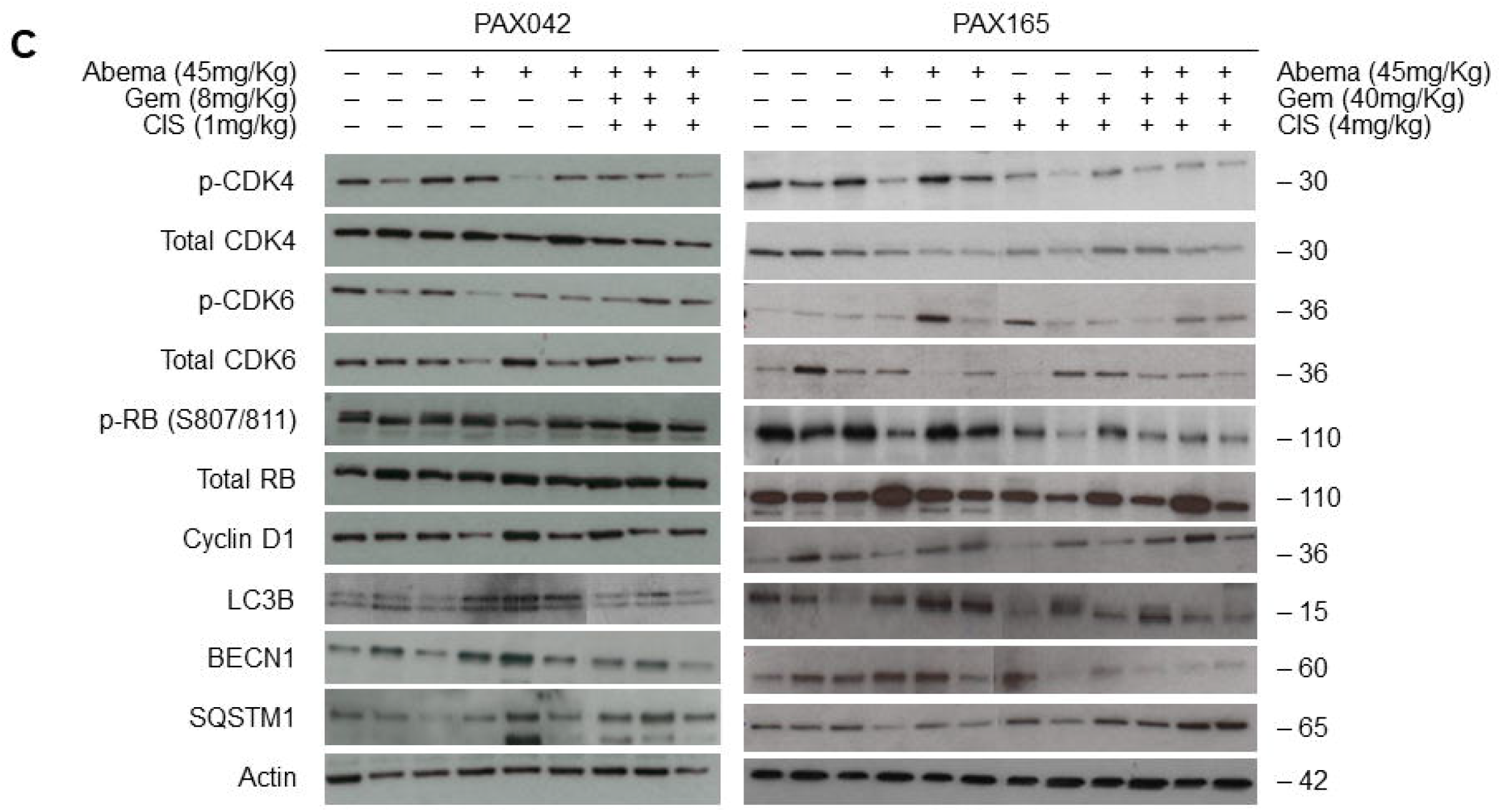
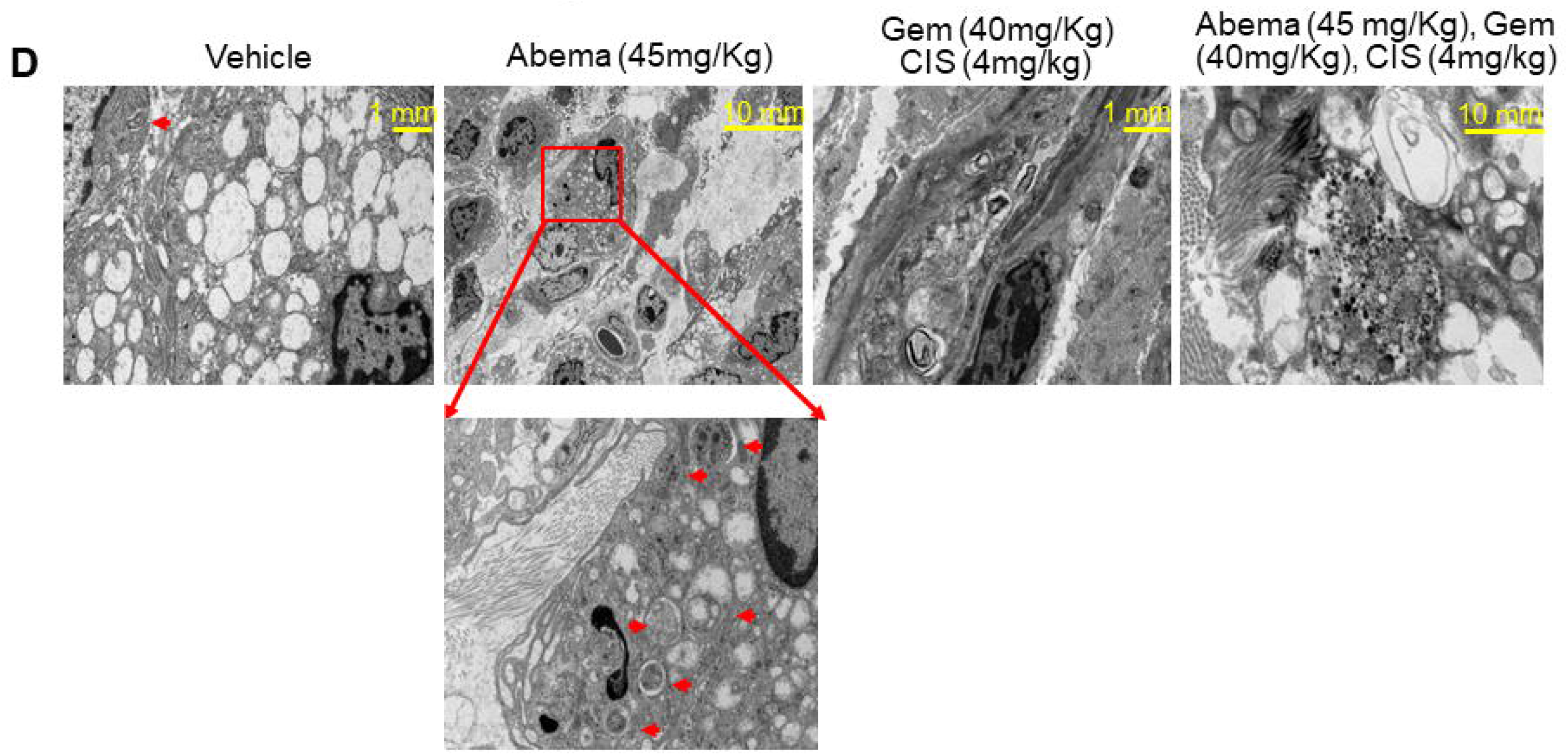
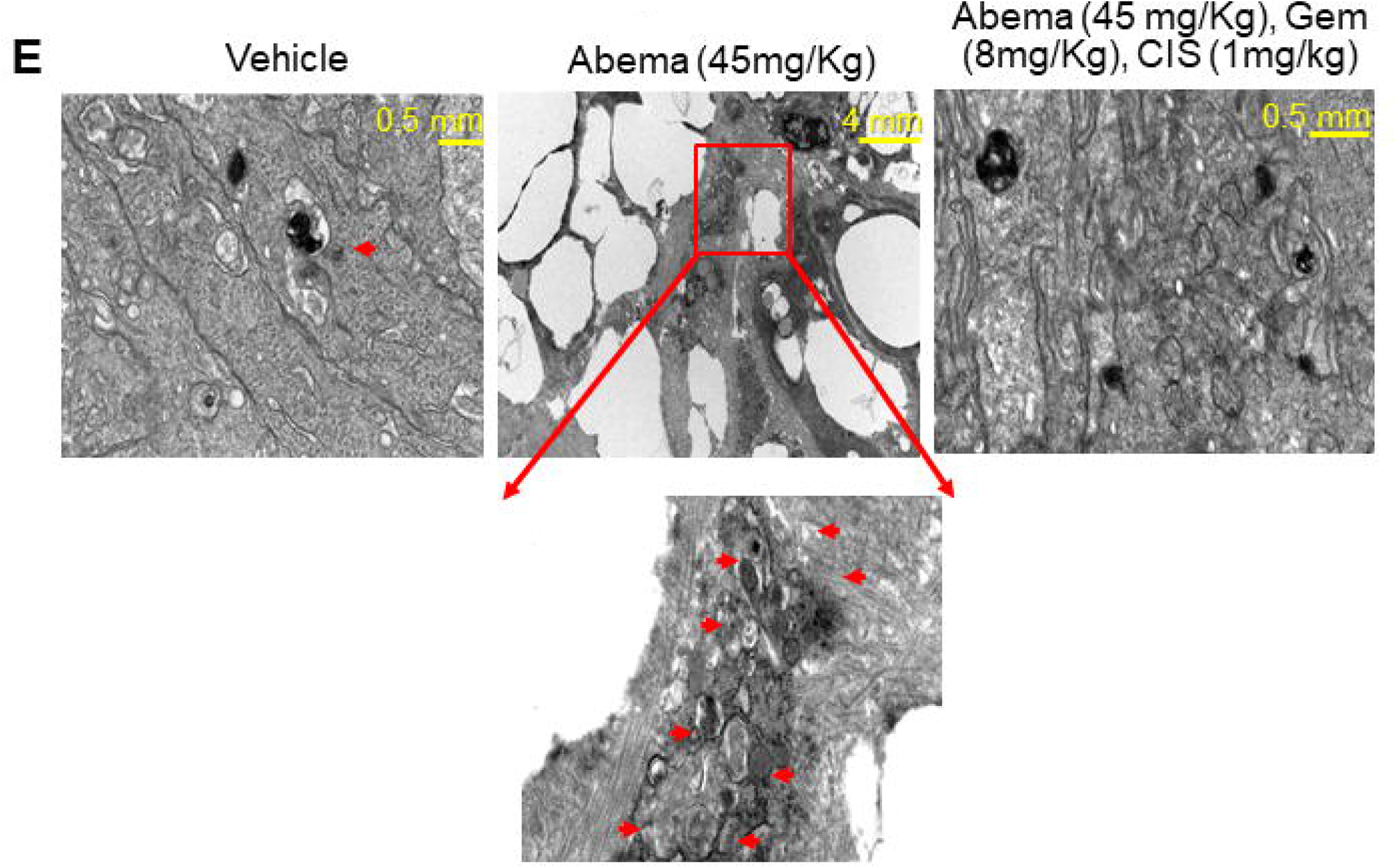
Impact of triplet combination of abemaciclib, gemcitabine and cisplatin on cellular effects of *in vivo* BTC models: Representative images for indicated assays in the end of study tumors in PAX165 (A) and PAX042 (B), the top panel shows histological evaluation of cellular proliferation via Ki67 staining (Ki67, Red; Nuclei, blue DAPI), the middle panel shows apoptotic cell death via TUNEL assay (TUNEL, Red; Nuclei, blue DAPI), and the bottom panel shows % necrosis via H & E staining, each for treatment cohort as labeled. All data represent mean ± s.e.m. (n=5) for each group from three independent experiments. (C) Western blot images of tumor lysates derived from three random mice per treatment cohort. Images of western blots were cropped to denote the relevant band(s) for clarity; b-actin was used as a loading control. (D, E) Representative TEM microphotographs of PAX165 (left) and PAX042 (right) treated with indicated concentrations of gemcitabine (Gem), cisplatin (CIS) and abemaciclib (Abema). Red boxes indicate magnified fields of regions containing autophagosomes or autolysosomes, indicated by red arrows. Error bars represent the standard error of mean values (N=5). (***P<0.0001, **P<0.001, *P<0.01).

Furthermore, we performed TUNEL staining on the tumor tissues as a surrogate measurement of apoptosis. The results revealed significantly increased % apoptosis in treated tumor tissues as compared to the control cohort. In PAX165, we observed 21% apoptotic activity in abemaciclib-treated (p=0.0116), 48% in doublet-treated (p<0.0001) and a 58% increase in triplet-treated (p<0.0001) as compared to the control cohort **(Figure 5A, middle panel)**. Congruently, in PAX042, results revealed a 22% increase in apoptotic activity in abemaciclib-treated (p=0.0023) in contrast to a 56% increase in triplet-treated cohort (p<0.0001) as compared to the control tumor tissues **(Figure 5B, middle panel)**.

Next, we investigated the histological differences in treated cohorts of both PDX models. H&E staining results revealed poorly differentiated tumors with slight therapy-related changes including cell enlargement, cytoplasmic vacuolization and cytoplasmic eosinophilia in treated cohorts, as compared to tumor tissue from the control cohorts. Compared to the control cohort mice, PAX165 H&E staining demonstrated 21%, 48% and 58% relative percent necrosis for abemaciclib-treated (p=0.01161), doublet-treated (p<0.0001) and triplet-treated (p<0.0001) cohorts respectively **(Figure 5A, bottom panel)**. In PAX042, relative percent necrosis for abemaciclib-treated was 22% (p=0.0023) and in triplet-treated was 56% (p<0.0001) as compared to controls **(Figure 5B, bottom panel)**. Taken together, these results suggest enhanced anti-tumor efficacy of the triplet-combination treatment as compared to the doublet or monotherapy abemaciclib treatment in *in vivo* CCA PDX models.

To further assess the functional and mechanistic impact of triplet-combination on the CCA PDX models, we evaluated the end of study tumor tissues (N=3) for pathway inhibition and other autophagy markers. The densitometry analyses of abemaciclib and triplet-treated cohort revealed a decrease in median protein expression of CDK4 (30.56 ± 0.22% and 47 ± 0.18%), CDK6 (16.50 ± 0.20% and 14.13 ± 0.49%), RB1 (20.73 ± 0.27% and 27.46 ± 0.46%) and CCND1 (12.23 ± 4.50% and 4.31 ± 2.49%) activity. In contrast, the doublet-treatment had a variable effect on CDK4 (33.21 ± 3.34%), CDK6 (no effect) or RB1 (26.92 ± 4.56%) activity **(Figure 5C and Supplementary Figure 6)**. This clearly indicates that triplet-combination treatment elicits *in vivo* anti-tumor activity in a cell-cycle dependent manner.

Finally, we investigated the effect of triplet-combination on key autophagy markers in the treated PDX tumor tissues. Abemaciclib treated cohorts of PAX165 and PAX042 demonstrated a significant increase in LC3B-II (72.51 ± 0.24%) and BECN1 (76.90 ± 0.63%) levels and decrease in SQSTM1 (30.01 ± 0.58%) protein levels as compared to the control cohort **(Figure 5C)**. Moreover, the triplet-combination treated cohorts of both the PDX models showed a significant decrease in LC3B-II (9.31 ± 1.31%) and BECN1 (41.50 ± 4.56%) protein levels **(Figure 5C)**. We also assessed these tumors for autophagic structures via transmission electron microscopy (TEM). TEM showed double-membrane autophagosomes and autolysomes in the abemaciclib treated cohort of PAX165 **(Figure 5D)** and PAX042 **(Figure 5E)**. Notably, TEM analysis of the triplet-treated cohort also revealed both autophagosomes and autolysosomes **(Figure 5D, E)**. However, due to the poorly differentiated histology, a quantitative comparison between the cohorts could not be performed.

Collectively, these results demonstrate that the combination of gemcitabine and cisplatin with abemaciclib mitigates their concomitant limitations i.e. toxicity and autophagy induction respectively, thus potentiating the efficacy of triplet-combination treatment in the CCA PDX models.

## Discussion

Despite ongoing advances, treatment options for patients with advanced BTC still remain limited. Combination of gemcitabine and cisplatin is the standard of care regimen that modestly improves ORR but has considerable toxicity and negative impacts on quality of life(3,46). Pathogenesis of BTC is driven by multiple actionable genomic events including alterations in cyclinD1-CDK4/6-CDKN2A-Rb pathway and loss of *CDKN2A*(5). Thus, BTCs could be classified as CDK4/6-dependent tumors. However, the molecular determinants of CDK4/6 sensitivity spectrum are not well understood and validation in clinical studies has not been fully undertaken(13). Herein we report that BTC cell lines are sensitive to the three FDA-approved CDK4/6 inhibitors with variable potencies but do not show any correlation with the underlying genomic alterations in the cyclinD1-CDK4/6-CDKN2A-RB pathway.

Development of therapeutic resistance is common and CDK4/6 inhibitors have been found to be more effective in combination with other agents that potentially circumvent these limitations (21,25,47). Several preclinical and clinical studies investigating this combination in non-BTC cancers have elucidated enhanced therapeutic efficacy associated with host protection and maintenance(41). However, the underlying mechanisms still need to be determined. Our findings demonstrate CDK4/6 inhibitor (abemaciclib) in combination with gemcitabine/cisplatin-chemotherapy regimen (gemcitabine and cisplatin) is synergistic *in vitro* and *in vivo* BTC models. Furthermore, we showed that the triplet-combination synergism is cell cycle dependent as there was a significant reduction of cyclinD1-CDK4/6-CDKN2A-Rb pathway activation, and an increase in cell cycle arrest and cell death as compared to the doublet and abemaciclib monotherapy. Although the inhibitory effect of triplet-combination on *RRM1* expression might suggest that synergism is cell-cycle independent, the increase in both cell cycle arrest and cell death clearly indicate that gemcitabine and cisplatin mutually cooperate with abemaciclib to strike a dynamic equipoise of cytostatic and cytotoxic response.

Progressively more studies have evaluated the role of autophagy as a mechanism of resistance to pharmacological CDK4/6 inhibition, specifically in breast and pancreatic cancer(48). These studies suggest that co-inhibition of CDK4/6 and autophagy might improve anti-tumor efficacy in cancers with an intact G1-S transition(21). Here, we used a suite of *in vitro* assays to demonstrate that triplet combination significantly reduces the autophagic flux as compared to doublet or abemaciclib monotherapy. Studies by Lefort *et al*. have shown an association of higher expression of autophagy markers like *LC3B* with tumor aggressiveness or residual disease post chemotherapy(49,50). In contrast, our findings suggest that gemcitabine/cisplatin chemotherapy did not have any effect on *LC3B* expression in our panel of BTC cell lines. Our results provide strong mechanistic evidence underlying cooperation of standard chemotherapy and CDK4/6 inhibitors.

Acquired resistance to CDK4/6 inhibitors has been linked to amplification of *CDK6* in preclinical models of several solid tumors(18,44). Here we report, insignificant anti-tumor activity to CDK4/6 inhibitor monotherapy in both PAX165 (*CDK6*^gain^) and PAX042 (*CDK6*^wt^) CCA PDX models, irrespective of their *CDK6* amplification status. Thus, CDK6 might not be a decisive resistance marker to CDK4/6 inhibitors in BTC. The doublet-combination of cytotoxic gemcitabine and cisplatin is highly efficacious in PAX165. Due to highly cytotoxic combination-effect of gemcitabine and cisplatin in PAX165, synergy-effect with abemaciclib could not be discerned in the triplet-combination cohort. However, our results clearly refute the alleged antagonism associated with the combination of chemotherapy with CDK4/6 inhibitors(25–29). Data from our pre-clinical assessment in *in vitro* and *in vivo* provides compelling evidence that the combination of CDK4/6 inhibitors with cytotoxic chemotherapy is based on synergistic mechanisms which mitigate chemotherapy-related toxicity, prevent abemaciclib resistance through avoidance of autophagy, and thereby enable a durable response as evidenced by survival benefit in pre-clinical BTC models. These promising preclinical results provide a strong mechanistic foundation for further clinical evaluation of the combination of gemcitabine/cisplatin with CDK4/6 inhibitors in advanced BTC patients.

## Abbreviations

BTC: biliary tract cancer;
SOC: standard of care;
CCA: cholangiocarcinoma;
GP: gemcitabine and cisplatin;
ORR: overall response rate;
CDK: cyclin dependent kinase;
CDKN2A: CDK inhibitor 2a;
RB: retinoblastoma;
HR: hormone receptor;
MBC: metastatic breast cancer;
PDAC: pancreatic ductal adenocarcinoma;
TNBC: triple-negative breast cancer;
NSCLC: non-small cell lung cancer;
RRM1: ribonucleotide reductase catalytic subunit M1;
DDR: drug dose response;
IC50: half-maximal inhibitory concentration;
CI: combination index;
PI: propidium iodide;
aCGH: array comparative genomic hybridization;
NOD: non-obese diabetic;
PDX: patient derived xenograft;
IP: intraperitoneal;
IHC: immunohistochemistry;
H&E: hematoxylin and eosin stain;
TUNEL: terminal deoxynucleotide transferase dUTP nick-end labeling;
R2: linear regression.

## Author Contributions

MA performed experiments, analyzed data and wrote the manuscript. JMB analyzed data and wrote the manuscript. AA, JY, RAR and JLL performed animal experiments and analyzed data. XC, PLSUJ, CRD, ATB, SIG, JBE, YZ, BMN, NM, ELE, MAS, HEK and KHB analyzed data. EB performed the targeted sequencing experiments and analyzed data. MTB conceived and directed the aCGH studies and analyzed data. MBS, ASM, LRR, TSB, DHA, MJT and MJB conceived and directed the study. All authors reviewed and approved the final manuscript.

