## Supplementary Methods for "Synergistic Combination of Cytotoxic Chemotherapy and Cyclin Dependent Kinase 4/6 Inhibitors in Biliary Tract Cancers"

**Supplementary Materials and Methods**

**Cell lines, drugs, antibodies and assay reagents**

Twenty one human biliary tract cancer cell lines were used in this study **(Table 1)** and all cultures maintained at 37°C in the presence of 5% CO_2_. For authentication, all BTC cell lines were profiled for using a targeted biliary cancer NGS sequencing panel. Palbociclib (#S1116), ribociclib (#S7440) and abemaciclib (#S5716) were obtained from SelleckChem LLC (Houston, TX). Clinical grade gemcitabine and cisplatin were obtained from the Mayo Clinic Cancer Center pharmacy. The APO-BrdU™ TUNEL Assay Kit (#A23210) was purchased from ThermoFisher Scientific (Waltham, MA USA). Alexa Fluor® 647 Anti-BrdU antibody (364107) was purchased from BioLegend. Ki-67 (D2H10) Rabbit mAb (IHC Specific; #9027) was purchased from Cell Signaling. Goat anti-rabbit Alexa-594 secondary rabbit (#R37117) was purchased from Invitrogen (Carlsbad, CA). Cell Titer-Glo® (#G7570) was obtained from Promega. Crystal violet was obtained from Sigma. FITC Annexin V Apoptosis Detection Kit I (RUO; #556547) was purchased from BD Biosciences. The Autophagy detection kit (ab139484) was obtained from Abcam (Cambridge, MA). Antibodies against phospho-CDK4 (Thr172; #5884)), total CDK4 (D9G3E; #12790), total CDK6 (D4S8S; #13331), phospho-RB (Ser807/811; #9308), total RB (4H1; #9309), cyclin D1 ((E3P5S; #55506), LC3A/B (#12741) and beclin1 (#3495) were purchased from Cell Signaling Technology (Danvers, MA). Anti-CDK6 (phosphor Y24; #ab131469) was purchased from Abcam (Cambridge, MA), anti-SQSTM1 (#sc-28359) purchased from Santa Cruz Biotechnology (Dallas, Texas) and Anti-β-Actin (#A00702-100) antibody was purchased from GenScript (Piscataway, NJ).

**Drug dose response studies**

Cells were plated in 384-well plates (Greiner Bio-One, NC USA) at optimized cell density and dosed 24 hours later (post-adherence). Relative luminescent signal was measured with CellTiter-Glo (CTG) after 144 hours drug exposure using a Cytation3 plate reader (BioTek, VT USA), and relative cell numbers (expressed as % viability) were calculated. Ten serially-diluted doses of palbociclib, ribociclib or abemaciclib, each in technical quadruplicates per biological replicate were evaluated. Prism version 7.03 (Prism Software Corporation, CA USA) was used to calculate concentrations corresponding to 50% viability (IC_50_). We evaluated eight doses of gemcitabine or cisplatin were combined with six doses of abemaciclib (54 dose combinations) in quadruplicate, analyzed and visualized with Combenefit software(1) using the Loewe model. After outlier removal, synergism of at least three doses of each drug was orthogonally confirmed in CalcuSyn analysis, version 2.1 (Biosoft, GB-UK).

**Cellular proliferation and death assays**

Cells were plated at 10e3 cells/well in a 24-well plate. After an adherence time of 24 hours, 1µM of palbociclib, ribociclib or abemaciclib was added. For combination analysis, abemaciclib [0.5µM] was added with or without doublet-gemcitabine [1nM] and cisplatin [1µM], and cells were incubated for 72 hours. Cells were then fixed with methanol-acetic acid (3:1) solution and stained with 0.5% crystal violet. The number of colonies, defined as ≥50 cells/colony, was counted manually by light microscopy. For cell cycle analysis, adherent cells were washed twice with ice-cold PBS, scraped off from the plate in 1 mL cold PBS, centrifuged at 1000rpm for 5 minutes before re-suspending in 500µl of NP-40 staining solution. NP40 staining solution contains 1% sodium citrate, 1mg/ml PI solution, 1% NP-40, 10mg/ml RNaseA and water. For cell death assays, cells were suspended in binding buffer at 10e3 cells/mL, Annexin V antibody added at [1:20] and propidium iodide was added [5 μg/mL]. After incubating for 15 minutes, cells were analyzed on LSR Fortessa (BD Biosciences, CA USA). The Autophagy Detection Kit (Abcam #ab139484) was used according to the manufacturers’ recommendations. Green detection reagent was added after 72 hours, and fluorescence was measured. DNA content flow sorting was performed on frozen PDX tissue samples as reported previously(2).

**Next generation sequencing and copy number analyses**

RNA sequencing, targeted panel sequencing and array comparative genomic analysis (aCGH) analysis were performed as described previously(2).

**Immunofluorescence**

For Lysotracker staining, cells were treated with abemaciclib or triplet combination in 35mm glass bottom cell imaging dishes. After 72 hours, 1µM lysotracker deep red was added per dish and incubated at 37°C. After 1 hour, probe containing media was replaced with fresh media and stained cells were visualized under the Cy5 signal by confocal microscope. For RFP-GFP-LC3 dual reporter assay, cells were transfected with ptf-LC3 vector using Lipofectamine 2000 as per manufacturer’s instructions. After 48 hours, these cells were treated with abemaciclib or triplet combination for 48 hours, washed with 1X PBS, fixed briefly (for 10 min) with 4% paraformaldehyde and mounted with Vectashield mounting media. They were then visualized with a Zeiss Confocal microscope LSM880 using the 488 and 643nm channels to image the GFP+ve and RFP+ve LC3 puncta respectively. The puncta were quantified using ImageJ to obtain the number of autophagosomes (yellow: RFP + GFP +) and autolysosomes (red: RFP +).

**Transmission electron microscopy**

Tissue samples were fixed in Trumps fixative (pH 7.2) at 4°C overnight, and then microwave processed using a Leica EM AMW microwave processor: Rinsed in phosphate buffer, post-fixed in 1% osmium tetroxide, stained with 2% uranyl acetate, dehydrated through a graded series of ethanols, and infiltrated with epoxy resin via acetone. They were then embedded into pure Spurr resin blocks. 100nm (or 0.1 μm) ultra-thin sections were cut and mounted on 200-mesh copper grids, post-stained with lead citrate, and observed under a JEOL (JEOL USA, Inc; MA, USA) JEM-1400 and JEM-1400 Plus transmission electron microscopes at 80kV.
