## Supplementary Tables for "Synergistic Combination of Cytotoxic Chemotherapy and Cyclin Dependent Kinase 4/6 Inhibitors in Biliary Tract Cancers"

**Table 1:** **Biliary Tract Cancer cell lines.** Panel of 21 biliary tract cancer cell lines used in the study. CCA – cholangiocarcinoma; eCCA – extrahepatic CCA; GBC – Gall bladder carcinoma; iCCA – intrahepatic CCA; FBS – fetal bovine serum; AA – antibiotic agent; Gift from Dr. Gores – Dr. Gregory Gores at Mayo Clinic, Rochester, MN; RIKEN - National Bio-Resource Project of the MEXT, Japan; KCLB - Korean cell line bank; JCRB - Japanese Collection of Research Bio resources; DSMZ - German Collection of Microorganisms and Cell Cultures.

| **Cell line** | **Site** | **Media** | **Source** |
| --- | --- | --- | --- |
| CAK-1 | eCCA | RPMI1640 + 10%FBS + AA | Gift from Dr. Gores |
| EGI-1 | eCCA | RPMI1640 + 10%FBS + AA | DSMZ |
| GBD-1 | GBC | RPMI1640 + 10%FBS + AA | Gift from Dr. Gores |
| H69 | "normal" cholangiocyte | DMEM/F12 + 10%FBS +GFs | Gift from Dr. Gores |
| HuCCT1 | iCCA | DMEM + 10%FBS + AA | JCRB |
| HuH-28 | iCCA | RPMI1640 + 10%FBS + AA | JCRB |
| KKU-100 | eCCA | DMEM + 10%FBS + AA | JCRB |
| KKU-213 | iCCA (Mixed Papillary and non-papillary) | DMEM + 10%FBS + AA | JCRB |
| KMCH-1 | HCC/CCA | DMEM + 10%FBS + AA | Gift from Dr. Gores |
| MMNK-1 | experimentally-immortalized cholangiocyte | DMEM + 10%FBS + AA | JCRB |
| RBE | iCCA | RPMI1640 + 10%FBS + AA | RIKEN |
| SNU-1079 | iCCA | RPMI1640 + 10%FBS + AA | KCLB |
| SNU-1196 | eCCA | RPMI1640 + 10%FBS + AA | KCLB |
| SNU-245 | eCCA | RPMI1640 + 10%FBS + AA | KCLB |
| SNU-308 | GBC | RPMI1640 + 10%FBS + AA | KCLB |
| SNU-478 | Ampulla of Vater | RPMI1640 + 10%FBS + AA | KCLB |
| SNU-869 | Ampulla of Vater | RPMI1640 + 10%FBS + AA | KCLB |
| SSP-25 | iCCA | RPMI1640 + 10%FBS + AA | RIKEN |
| TFK-1 | eCCA | RPMI1640 + 10%FBS + AA | RIKEN |
| TKKK | iCCA | DMEM + 10%FBS + AA | RIKEN |
| YSCCC | iCCA | DMEM + 10%FBS + AA | RIKEN |

**Table 2:** **Comparison of *in vitro* drug dose response for monotherapy palbociclib, ribociclib and abemaciclib in BTC cell lines.** Mean half-maximal inhibitory concentration (IC_50_) values for palbociclib, ribociclib and abemaciclib from three biological replicate experiments shown with standard deviations. The indicated delta represents the fold difference between the highest and lowest IC50 values for palbociclib, ribociclib and abemaciclib.

| **Cell Line** | **Palbociclib IC_50_ [µM]** | **Ribociclib IC_50_ [µM]** | **Abemaciclib IC_50_ [µM]** |
| --- | --- | --- | --- |
| CAK-1 | 7.76 | 15.77 | 1.18 |
| TFK-1 | 7.01 | 19.8 | 1.02 |
| SNU-308 | 3.08 | 9.19 | 1.20 |
| GBD-1 | 7.87 | 14.57 | 0.99 |
| HuCCT1 | 2.48 | 10.60 | 0.99 |
| EGI-1 | 0.50 | 5.01 | 0.29 |
| HuH-28 | 7.25 | 15.31 | 0.56 |
| KMCH-1 | 1.29 | 3.41 | 0.12 |
| KKU-100 | 5.01 | 15.00 | 0.19 |
| MMNK-1 | 2.48 | 17.73 | 0.81 |
| KKU-213 | 6.58 | 19.52 | 1.38 |
| SNU-245 | 7.13 | 15.8 | 0.02 |
| SNU-478 | 2.13 | 6.57 | 0.46 |
| SNU-869 | 7.71 | 13.52 | 0.29 |
| SNU-1079 | 7.57 | 7.57 | 0.28 |
| SNU-1196 | 6.66 | 16.01 | 0.49 |
| RBE | 4.04 | 1.88 | 0.01 |
| YSCCC | 7.88 | 2.46 | 0.02 |
| SSP-25 | 5.40 | 13.20 | 0.54 |
| TKKK | 6.20 | 11.00 | 0.78 |
| H69 | 8.24 | 22.18 | 5.17 |
| **Median** | **6.58** | **13.52** | **0.54** |
| **STDEV** | **2.45** | **5.80** | **1.06** |
| **Delta** | **2.69** | **2.33** | **0.51** |
