## Supplementary Figures for "Synergistic Combination of Cytotoxic Chemotherapy and Cyclin Dependent Kinase 4/6 Inhibitors in Biliary Tract Cancers"

### Slide 1
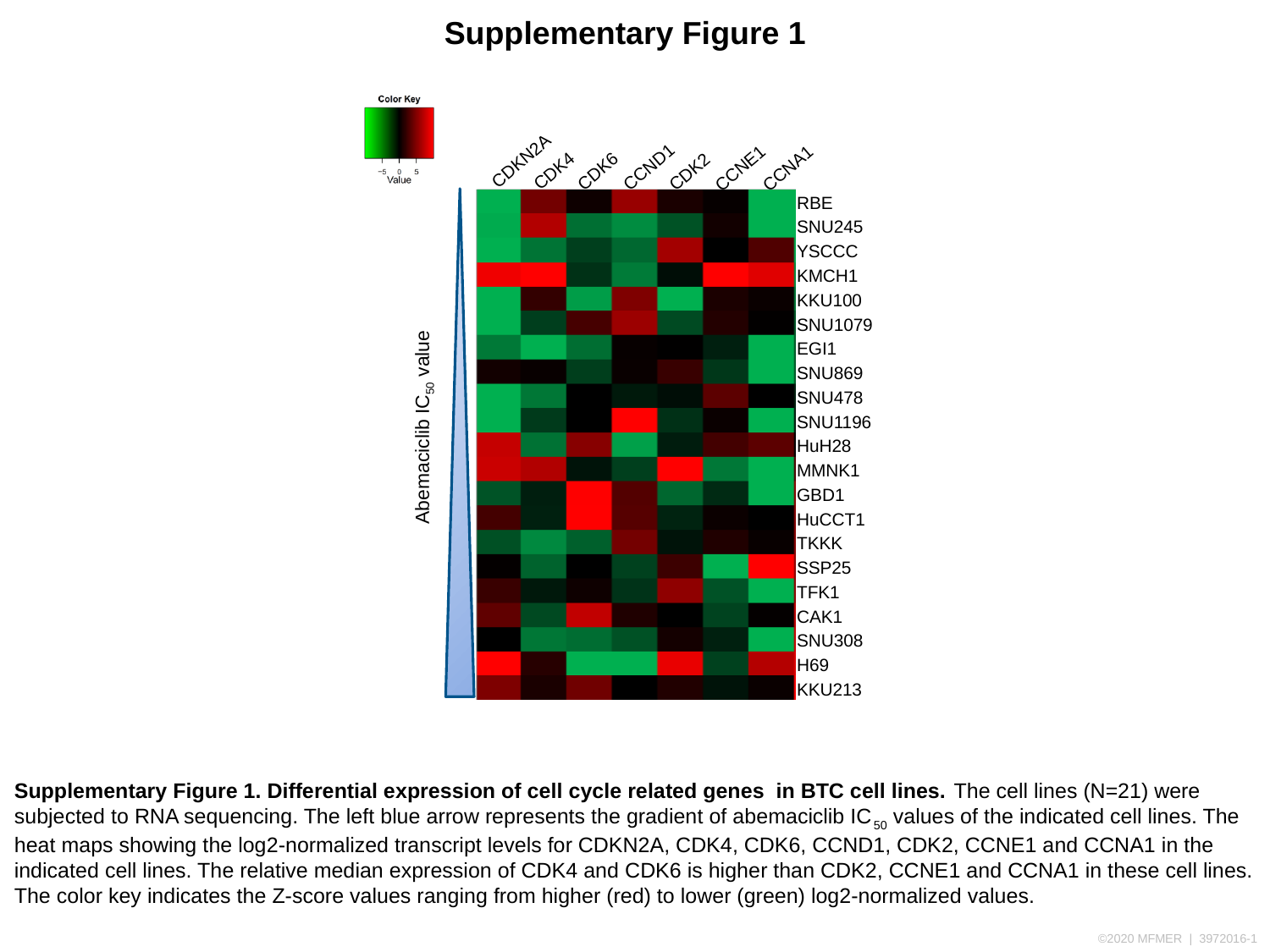

Supplementary Figure 1
CDKN2A
CDK4
CDK6
CCND1
CDK2
CCNE1
CCNA1
| RBE |
| --- |
| SNU245 |
| YSCCC |
| KMCH1 |
| KKU100 |
| SNU1079 |
| EGI1 |
| SNU869 |
| SNU478 |
| SNU1196 |
| HuH28 |
| MMNK1 |
| GBD1 |
| HuCCT1 |
| TKKK |
| SSP25 |
| TFK1 |
| CAK1 |
| SNU308 |
| H69 |
| KKU213 |
Abemaciclib IC_50_ value
Supplementary Figure 1. Differential expression of cell cycle related genes in BTC cell lines. The cell lines (N=21) were subjected to RNA sequencing. The left blue arrow represents the gradient of abemaciclib IC_50_ values of the indicated cell lines. The heat maps showing the log2-normalized transcript levels for CDKN2A, CDK4, CDK6, CCND1, CDK2, CCNE1 and CCNA1 in the indicated cell lines. The relative median expression of CDK4 and CDK6 is higher than CDK2, CCNE1 and CCNA1 in these cell lines. The color key indicates the Z-score values ranging from higher (red) to lower (green) log2-normalized values.

### Slide 2
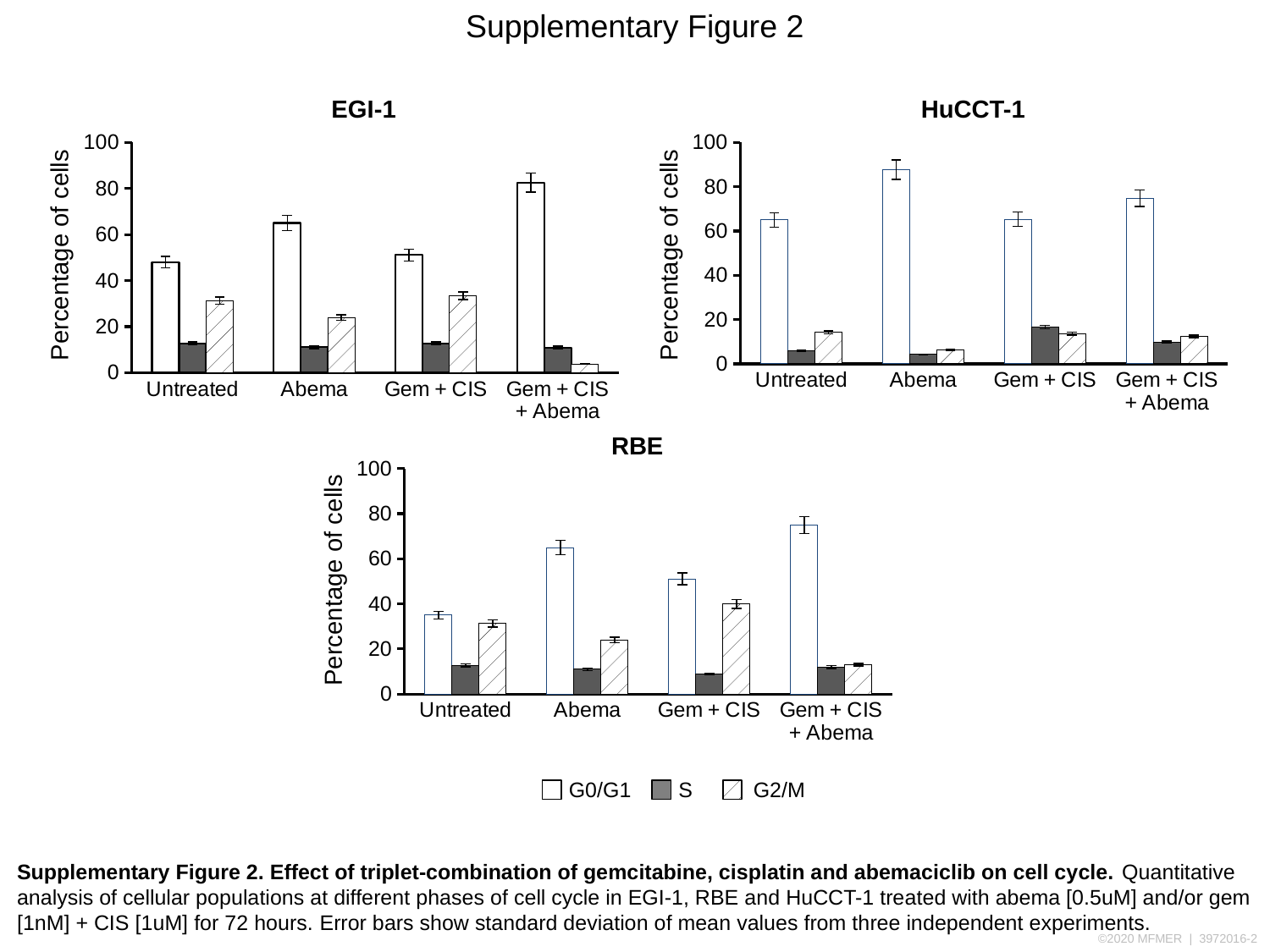

Supplementary Figure 2
EGI-1
HuCCT-1
#### Chart
| Category | G0/G1 | S | G2/M |
|---|---|---|---|
| Untreated | 48.0 | 12.7 | 31.3 |
| Abema | 65.0 | 11.0 | 24.0 |
| Gem + CIS | 51.1 | 12.7 | 33.4 |
| Gem + CIS + Abema | 82.5 | 10.9 | 3.8 |
#### Chart
| Category | G0/G1 | S | G2/M |
|---|---|---|---|
| Untreated | 65.0 | 5.89 | 14.2 |
| Abema | 87.6 | 4.25 | 6.28 |
| Gem + CIS | 65.3 | 16.6 | 13.7 |
| Gem + CIS + Abema | 74.7 | 9.76 | 12.4 |Percentage of cells
Percentage of cells
RBE
#### Chart
| Category | G0/G1 | S | G2/M |
|---|---|---|---|
| Untreated | 35.0 | 12.7 | 31.3 |
| Abema | 65.0 | 11.0 | 24.0 |
| Gem + CIS | 51.1 | 8.900000000000006 | 40.0 |
| Gem + CIS + Abema | 75.0 | 12.0 | 13.0 |Percentage of cells
G0/G1
S
G2/M
Supplementary Figure 2. Effect of triplet-combination of gemcitabine, cisplatin and abemaciclib on cell cycle. Quantitative analysis of cellular populations at different phases of cell cycle in EGI-1, RBE and HuCCT-1 treated with abema [0.5uM] and/or gem [1nM] + CIS [1uM] for 72 hours. Error bars show standard deviation of mean values from three independent experiments.

### Slide 3
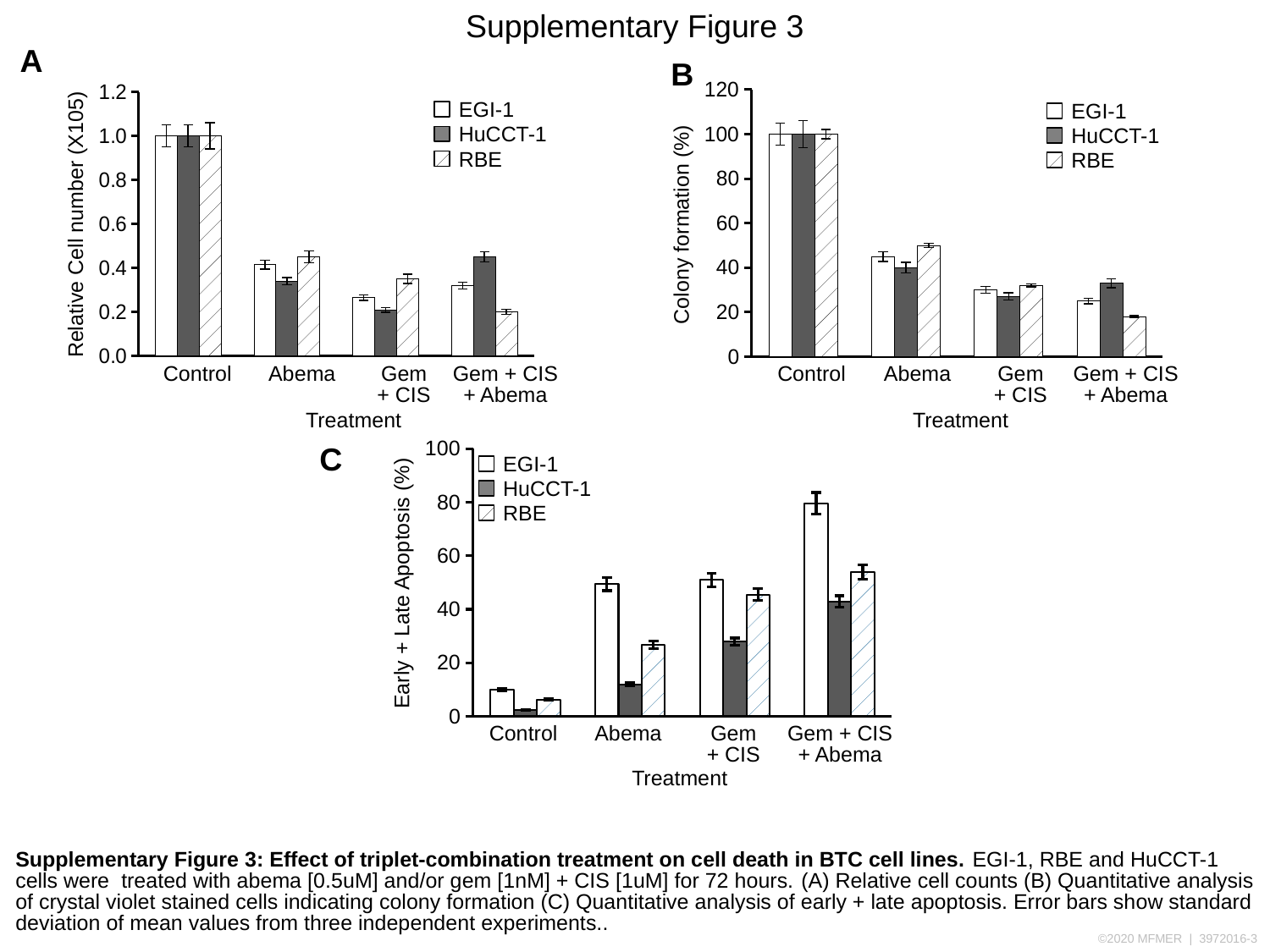

Supplementary Figure 3
A
B
#### Chart
| Category | EGI-1 | HuCCT-1 | RBE |
|---|---|---|---|
| Control | 1.0 | 1.0 | 1.0 |
| Abema | 0.41478734311210796 | 0.34 | 0.45 |
| Gem + CIS | 0.26525203847847895 | 0.21 | 0.35 |
| Gem + CIS + Abema | 0.31996061840941303 | 0.44999999999999996 | 0.19999999999999996 |
#### Chart
| Category | EGI-1 | HuCCT-1 | RBE |
|---|---|---|---|
| Control | 100.0 | 100.0 | 100.0 |
| Abema | 45.0 | 40.0 | 50.0 |
| Gem+CIS | 30.0 | 27.0 | 32.0 |
| Gem+CIS+Abema | 25.0 | 33.0 | 18.0 |EGI-1
HuCCT-1
RBE
EGI-1
HuCCT-1
RBE
Relative Cell number (X105)
Colony formation (%)
Control
Abema
Gem
+ CIS
Gem + CIS
+ Abema
Control
Abema
Gem
+ CIS
Gem + CIS
+ Abema
Treatment
Treatment
C
#### Chart
| Category | EGI-1 | HuCCT-1 | RBE |
|---|---|---|---|
| Untreated | 9.86 | 2.42 | 6.24 |
| Abema | 49.4 | 11.93 | 26.7 |
| Gem + CIS | 51.0 | 27.82 | 45.5 |
| Gem + CIS + Abema | 79.7 | 42.9 | 53.9 |EGI-1
HuCCT-1
RBE
Early + Late Apoptosis (%)
Control
Abema
Gem
+ CIS
Gem + CIS
+ Abema
Treatment
Supplementary Figure 3: Effect of triplet-combination treatment on cell death in BTC cell lines. EGI-1, RBE and HuCCT-1 cells were treated with abema [0.5uM] and/or gem [1nM] + CIS [1uM] for 72 hours. (A) Relative cell counts (B) Quantitative analysis of crystal violet stained cells indicating colony formation (C) Quantitative analysis of early + late apoptosis. Error bars show standard deviation of mean values from three independent experiments..

### Slide 4
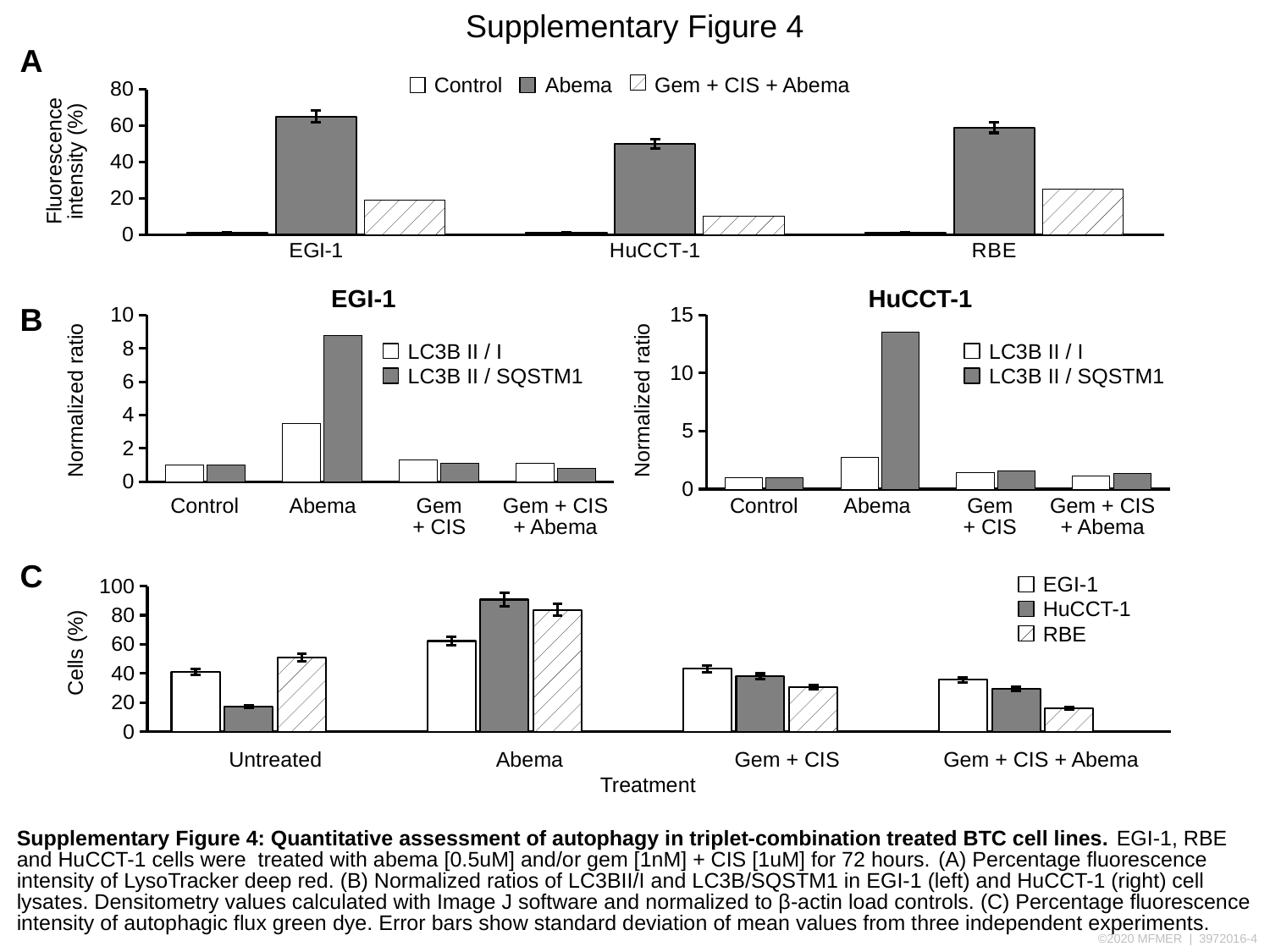

Supplementary Figure 4
A
Control
Abema
Gem + CIS + Abema
#### Chart
| Category | Control | Abema | Gem + CIS + Abema |
|---|---|---|---|
| EGI-1 | 1.0 | 65.0 | 19.0 |
| HuCCT-1 | 1.0 | 50.0 | 10.0 |
| RBE | 1.0 | 59.0 | 25.0 |Fluorescence
intensity (%)
EGI-1
HuCCT-1
B
#### Chart
| Category | LC3B II / I | LC3B II / SQSTM1 |
|---|---|---|
| Control | 1.0 | 1.0 |
| Abema | 3.5 | 8.75 |
| Gem + CIS | 1.3 | 1.0833333333333335 |
| Gem + CIS + Abema | 1.1 | 0.7857142857142858 |
#### Chart
| Category | LC3B II / I | LC3B II / SQSTM1 |
|---|---|---|
| Control | 1.0 | 1.0 |
| Abema | 2.7 | 13.5 |
| Gem + CIS | 1.4 | 1.5555555555555554 |
| Gem + CIS + Abema | 1.1 | 1.375 |LC3B II / I
LC3B II / SQSTM1
LC3B II / I
LC3B II / SQSTM1
Normalized ratio
Normalized ratio
Control
Abema
Gem
+ CIS
Gem + CIS
+ Abema
Control
Abema
Gem
+ CIS
Gem + CIS
+ Abema
C
EGI-1
HuCCT-1
RBE
#### Chart
| Category | EGI-1 | HuCCT-1 | RBE | Column1 |
|---|---|---|---|---|
| Untreated | 41.3 | 17.4 | 50.8 | None |
| Abema | 62.3 | 90.8 | 83.7 | None |
| Gem + CIS | 43.2 | 38.0 | 30.7 | None |
| Gem + CIS + Abema | 35.8 | 29.5 | 16.1 | None |
Cells (%)
Untreated
Abema
Gem + CIS
Gem + CIS + Abema
Treatment
Supplementary Figure 4: Quantitative assessment of autophagy in triplet-combination treated BTC cell lines. EGI-1, RBE and HuCCT-1 cells were treated with abema [0.5uM] and/or gem [1nM] + CIS [1uM] for 72 hours. (A) Percentage fluorescence intensity of LysoTracker deep red. (B) Normalized ratios of LC3BII/I and LC3B/SQSTM1 in EGI-1 (left) and HuCCT-1 (right) cell lysates. Densitometry values calculated with Image J software and normalized to β-actin load controls. (C) Percentage fluorescence intensity of autophagic flux green dye. Error bars show standard deviation of mean values from three independent experiments.

### Slide 5
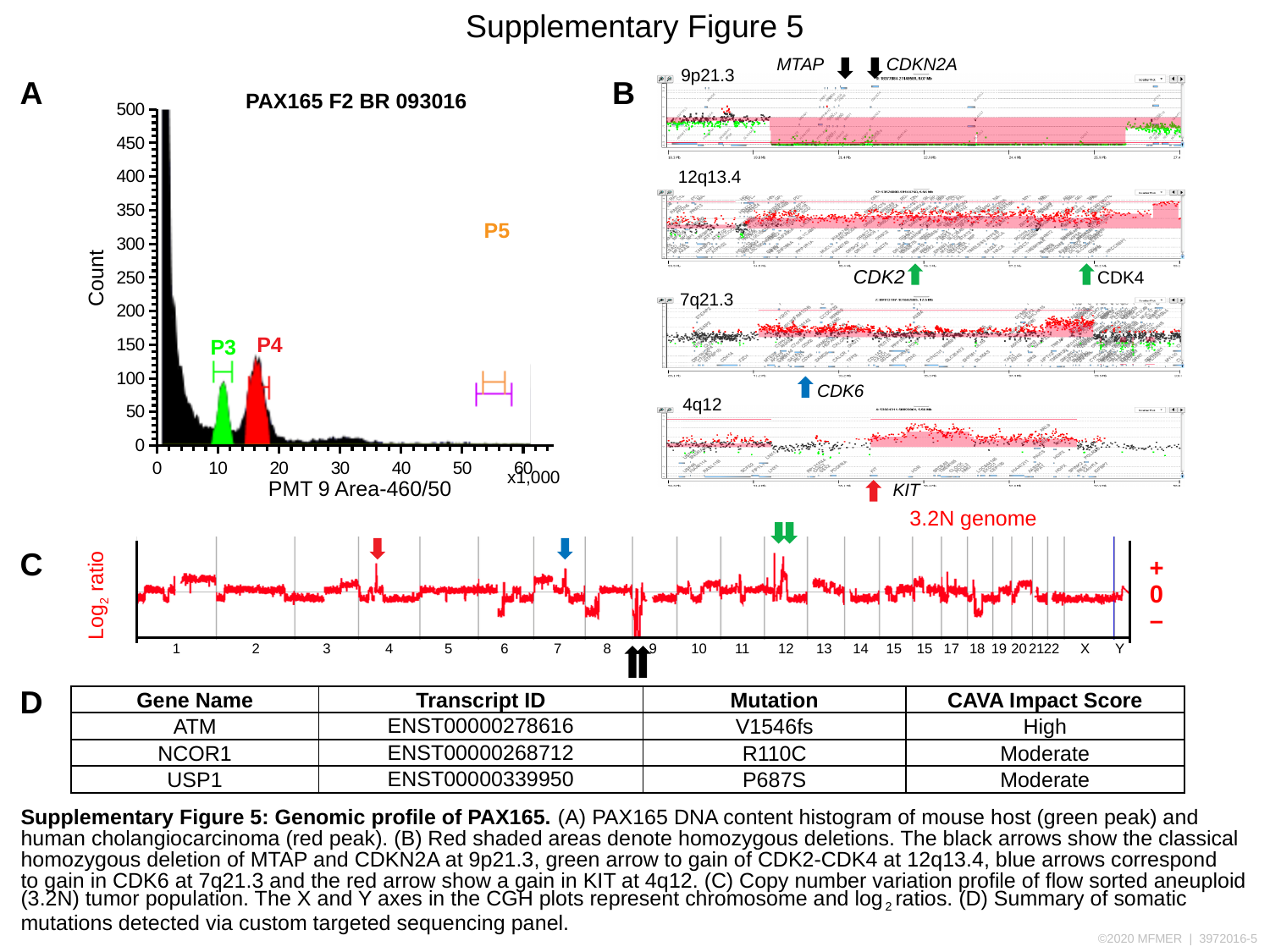

Supplementary Figure 5
MTAP
CDKN2A
9p21.3
A
B
PAX165 F2 BR 093016
#### Chart
| Category | Column1 |
|---|---|
12q13.4
P5
CDK2
Count
CDK4
7q21.3
P4
P3
CDK6
4q12
x1,000
PMT 9 Area-460/50
KIT
3.2N genome
C
+
0
Log_2_ ratio
–
1
2
3
4
5
6
7
8
9
10
11
12
13
14
15
15
17
18
19
20
21
22
X
Y
D
| Gene Name | Transcript ID | Mutation | CAVA Impact Score |
| --- | --- | --- | --- |
| ATM | ENST00000278616 | V1546fs | High |
| NCOR1 | ENST00000268712 | R110C | Moderate |
| USP1 | ENST00000339950 | P687S | Moderate |
Supplementary Figure 5: Genomic profile of PAX165. (A) PAX165 DNA content histogram of mouse host (green peak) and human cholangiocarcinoma (red peak). (B) Red shaded areas denote homozygous deletions. The black arrows show the classical homozygous deletion of MTAP and CDKN2A at 9p21.3, green arrow to gain of CDK2-CDK4 at 12q13.4, blue arrows correspond
to gain in CDK6 at 7q21.3 and the red arrow show a gain in KIT at 4q12. (C) Copy number variation profile of flow sorted aneuploid (3.2N) tumor population. The X and Y axes in the CGH plots represent chromosome and log_2_ ratios. (D) Summary of somatic mutations detected via custom targeted sequencing panel.

### Slide 6
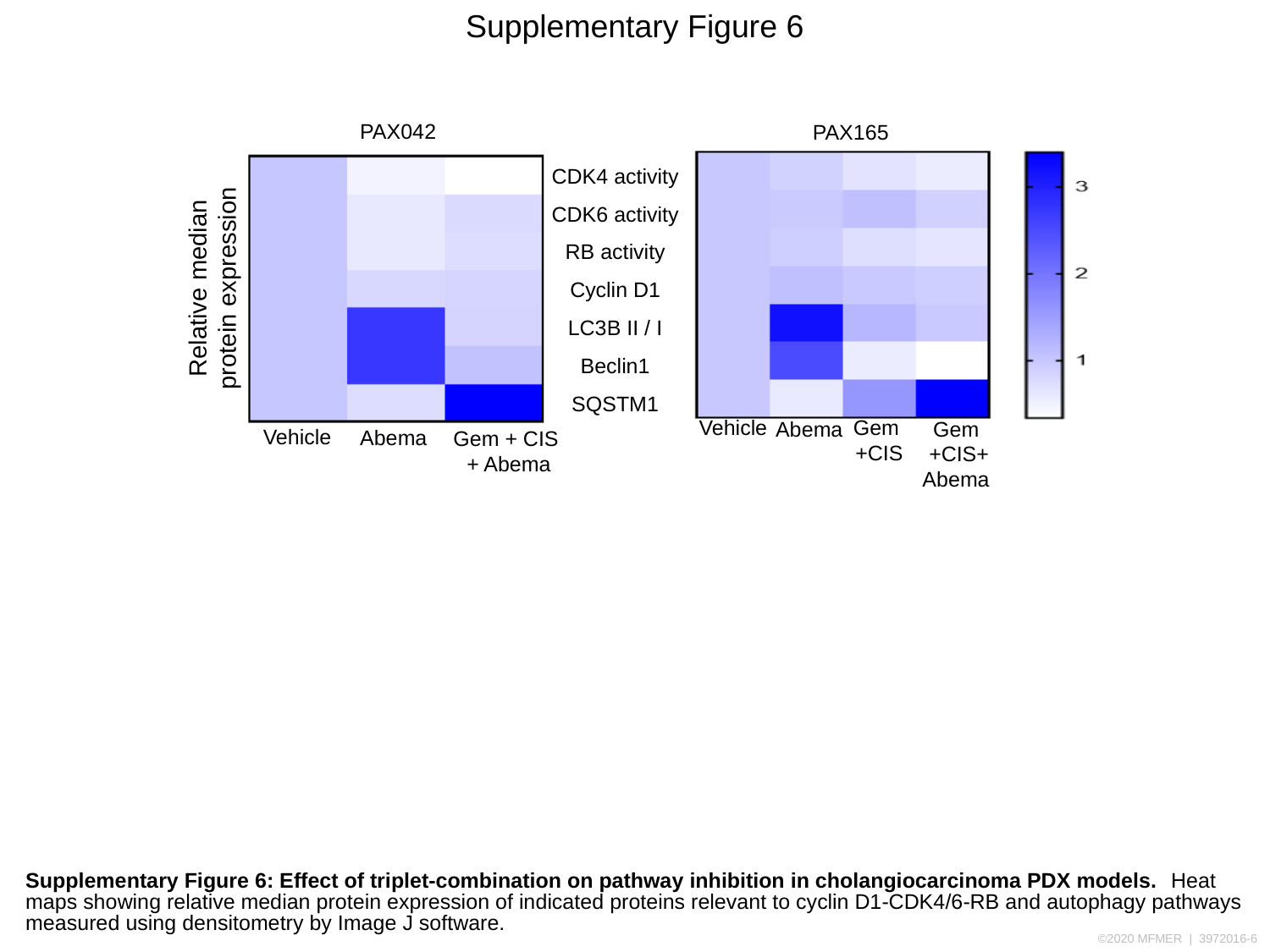

Supplementary Figure 6
PAX042
PAX165
CDK4 activity
CDK6 activity
RB activity
Cyclin D1
LC3B II / I
Beclin1
SQSTM1
Relative median
protein expression
Vehicle
Gem
+CIS
Abema
Gem
+CIS+
Abema
Vehicle
Abema
Gem + CIS
 + Abema
Supplementary Figure 6: Effect of triplet-combination on pathway inhibition in cholangiocarcinoma PDX models. Heat maps showing relative median protein expression of indicated proteins relevant to cyclin D1-CDK4/6-RB and autophagy pathways measured using densitometry by Image J software.
